# Ribosome stalling caused by the Argonaute-microRNA-SGS3 complex regulates the production of secondary siRNAs in plants

**DOI:** 10.1101/2020.09.10.288902

**Authors:** Hiro-oki Iwakawa, Andy Y.W. Lam, Akira Mine, Tomoya Fujita, Kaori Kiyokawa, Manabu Yoshikawa, Atsushi Takeda, Shintaro Iwasaki, Yukihide Tomari

## Abstract

The path of ribosomes on mRNAs can be impeded by various obstacles. One such example is halting of ribosome movement by microRNAs, though the exact mechanism and physiological role remain unclear. Here, we find that ribosome stalling caused by the Argonaute-microRNA-SGS3 complex regulates the production of secondary small interfering RNAs (siRNAs) in plants. We show that the double-stranded RNA-binding protein SGS3 directly interacts with the 3′ end of the microRNA in an Argonaute protein, resulting in ribosome stalling. Importantly, microRNA-mediated ribosome stalling positively correlates with efficient production of secondary siRNAs from target mRNAs. Our results illustrate a role for paused ribosomes in regulation of small RNA function that may have broad biological implications across the plant kingdom.

## Main

Ribosome movement can be interrupted by various factors including rare codons, special RNA structures and specific amino acid sequences called ribosome arrest peptides (Ito and Chiba, 2013; Schuller and Green, 2018). Although the physiological roles of such impediments are unclear, growing evidence indicates that ribosome stalling has diverse functions, including ER stress response, monitoring protein secretion, feedback regulation of methionine biosynthesis, quality control of mRNAs, and folding of nascent peptide chains (Ito and Chiba, 2013; Inada, 2017; Stein et al., 2019).

microRNAs (miRNAs) can cause ribosome stalling as well as inhibition of translation initiation, target RNA degradation or cleavage (Fabian et al., 2010; Iwakawa and Tomari, 2013; Iwakawa and Tomari, 2015; Hou et al., 2016; Li et al., 2016; Bazin et al., 2017; Zhang et al., 2018). To pause ribosomes, miRNAs need to form RNA-induced silencing complexes (RISCs) with Argonaute (AGO) protein, and extensively base-pair within the coding sequence (CDS) of the target mRNA (Iwakawa and Tomari, 2013; Hou et al., 2016; Zhang et al., 2018). However, these requirements are not sufficient for ribosome stalling in plants; although many plant miRNAs have their cleavable targets with perfect or near perfect complementary binding sites in CDS, only a few miRNA binding sites can induce ribosome stalling *in vivo* (Hou et al., 2016). Thus, unknown elements other than RISC binding should be required for miRNA-mediated ribosome pausing.

The biological function of the miRNA-mediated ribosome stalling also remains unclear. One plausible role of the miRNA-mediated ribosome stalling is inhibition of functional protein synthesis (Iwakawa and Tomari, 2013; Iwakawa and Tomari, 2015; Zhang et al., 2018). However, given the diverse functions of stalled ribosomes as mentioned above, miRNA-mediated ribosome pausing may have a role other than translation repression.

Here, we show that a dsRNA binding protein, SGS3, is a key determinant of miRNA-mediated ribosome stalling. SGS3 forms a complex on dsRNA protruding from the miR390-AGO7-target complex. These mechanisms also operate in the context of a distinct 22-nucleotide miRNA-AGO1-RISC complex. Importantly, we find that SGS3 and miRNA-mediated ribosome stalling positively correlates with efficient amplification of RNA silencing, suggesting a new role of ribosome pausing beyond inhibition of protein synthesis.

## Results

### The dsRNA-binding protein SGS3 is a specific enhancer for microRNA-mediated ribosome stalling

We first sought to find the miRNA-mediated ribosome stalling positions. To do this, we performed ribosome profiling, an approach that is based on sequencing of ribosome-protected footprints after RNase treatment (Ingolia et al., 2009), in *Arabidopsis* seedlings. Our data represented a 3-nucleotide periodicity along the ORF, a hallmark of translation elongation (Figure S1A). We combined this high-resolution ribosome profiling and the miRNA target prediction (Dai et al., 2018) to identify the ribosome-stalling position upstream of the predicted miRNA binding sites (Table S1). Along with earlier studies (Hou et al., 2016; Li et al., 2016; Bazin et al., 2017), our ribosome profiling has shown that specific miRNAs, including miR390 and miR173, can induce ribosome stalling 12– 13 nucleotide upstream of their binding sites in *Arabidopsis thaliana* Figure 1A–C and S1B–D). These particular miRNAs are known to trigger the production of phased secondary small interfering RNAs (siRNAs), called trans-acting siRNAs (tasiRNAs), from precursors called TAS RNAs (Liu et al., 2020). tasiRNA production requires various factors including AGO7 and AGO1, which form specific RISCs with miR390 and miR173, respectively (Montgomery et al., 2008; Endo et al., 2013; Liu et al., 2020). One important factor for tasiRNA biogenesis is SUPPRESSOR OF GENE SILENCING 3 (SGS3) (Mourrain et al., 2000; Peragine et al., 2004; Vazquez et al., 2004; Allen et al., 2005). Given that SGS3 forms cytoplasmic foci named “siRNA bodies” with AGO7 (Jouannet et al., 2012) and interacts with AGO1 associating with miR173 and other 22- nt small RNAs (Chen et al., 2010; Cuperus et al., 2010; Yoshikawa et al., 2013), we reasoned that SGS3 influences miRNA-mediated ribosome stalling. To test this idea, we examined the impact of an SGS3 mutation on miRNA-mediated ribosome stalling by comparing ribosome profiling in wild-type and *sgs3-11 Arabidopsis* seedlings (Peragine et al., 2004). We observed dramatic decreases in ribosome stalling in *sgs3-11* mutants (Figure 1B and C and S1B–D and S2). This reduction cannot be explained by a change in mRNA or miRNA abundance in the mutant (Figure 1B and C and S1–3). Thus, we concluded that SGS3 is required for ribosome stalling by miR390 and miR173. Given that *sgs3-11* mutation did not cause an overall decrease in ribosome occupancy (Figure S2), SGS3 is not a general ribosome stalling factor, but rather a specific stalling enhancer for miRNA-mediated ribosome stalling.

**Figure 1.**
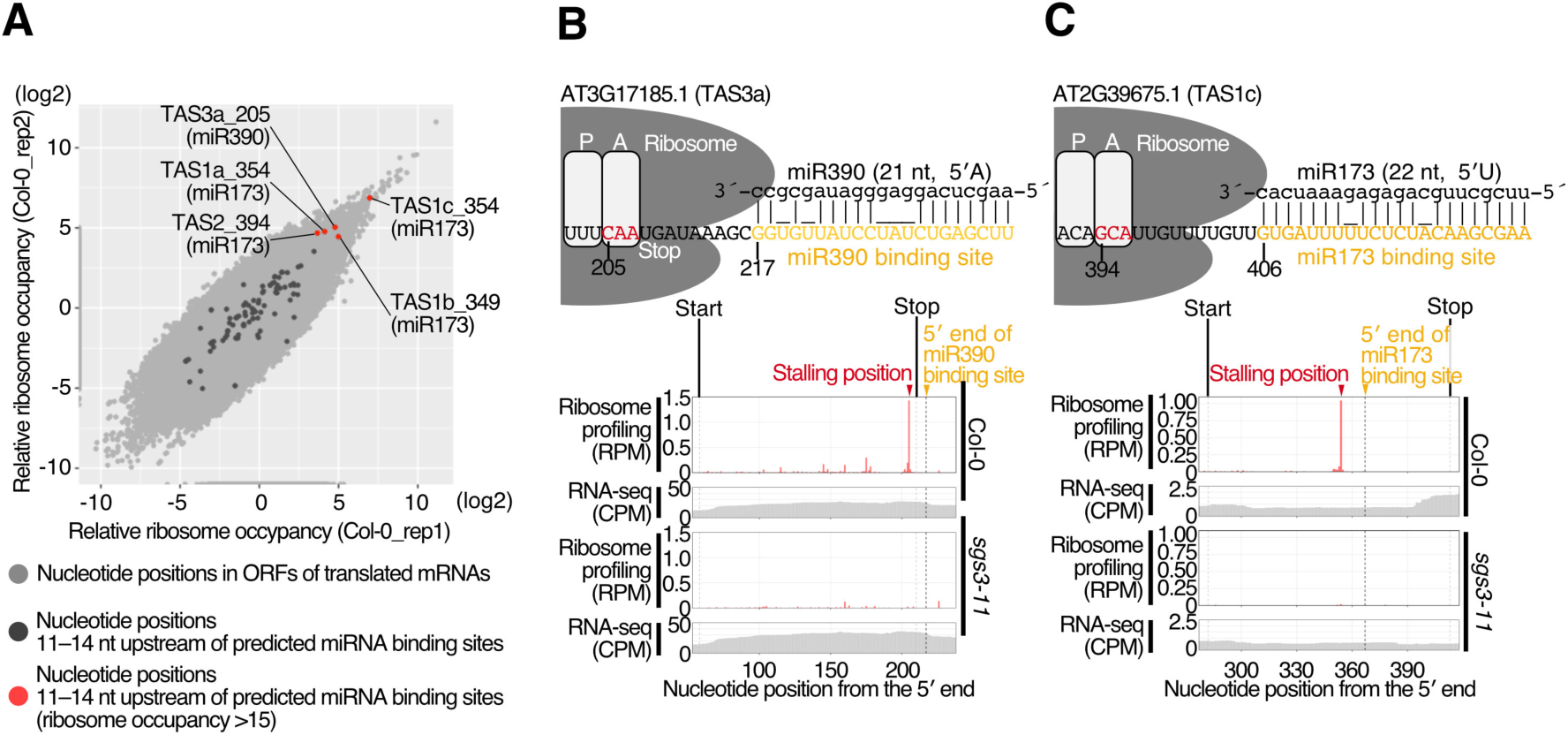
The dsRNA-binding protein SGS3 promotes microRNA-mediated ribosome stalling. (A) Scatter plot showing correlation of relative ribosome occupancy (Materials and Methods) between replicates (Col-0_rep1 and 2). The nucleotide positions with ribosome footprints (reads per million (RPM) over 0.05) in translating ORFs are shown in light gray. The nucleotide positions 11–14 nucleotide upstream of predicted miRNA binding sites are shown in dark gray (Table S1). In such positions, those with relative ribosome occupancy over 15 are shown in red (Table S1). (B, C) Ribosome footprints (A-site position) in RPM and RNA-seq in coverage per million (CPM) in wild-type or *sgs3-11* mutant seedlings are shown for the following transcripts: (B) AT3G17185.1 (TAS3a), a precursor of trans acting siRNAs (tasiRNAs) with miR390 binding sites; (C) AT2G39675.1 (TAS1c), a precursor of tasiRNAs with a miR173 binding site. See also Figure S1 and Table S1.

### SGS3 and RISC cooperatively stall ribosomes *in vitro*

Although our ribosome profiling data demonstrate the involvement of SGS3 and miRNAs in ribosome stalling, how these factors coordinately pause ribosomes was unclear. To reveal the mechanisms of miRNA- and SGS3-dependent ribosome stalling, we adopted a tobacco BY-2 cell-free system, which can recapitulate miRNA-mediated RNA silencing *in vitro* (Figure 2A) (Iki et al., 2010; Iwakawa and Tomari, 2013). We used TAS3a as a representative target RNA. TAS3a contains a short (51 codon) ORF and two miR390- binding sites: one is adjacent to the stop codon and immediately downstream of the ribosome stalling site, and the other is located well downstream of those elements (Figure 2B) (Axtell et al., 2006). Ribosome stalling within the short ORF was monitored by detecting peptidyl-tRNAs, a hallmark of ribosome stalling (Nakatogawa and Ito, 2001; Muto et al., 2006), by western blotting to a FLAG-tag inserted in the F-TAS3 ORF (Figure 2A and C). Western blotting followed a neutral pH gel electrophoresis that prevents hydrolysis of the ester linkage between the tRNA and amino acid (Nakatogawa and Ito, 2001), thus enabling us to detect peptidyl-tRNAs within stalled ribosomes through an ∼18 kDa upshift —the size of the tRNA moiety (Figure 2C). Translation of the reporter (F-TAS3) in the presence of AGO7-RISC led to a clear band-shift (Figure 2D and E). Disappearance of this signal after RNase treatment confirmed that the upshifted band corresponds to peptidyl-tRNA (Figure 2D).

**Figure 2.**
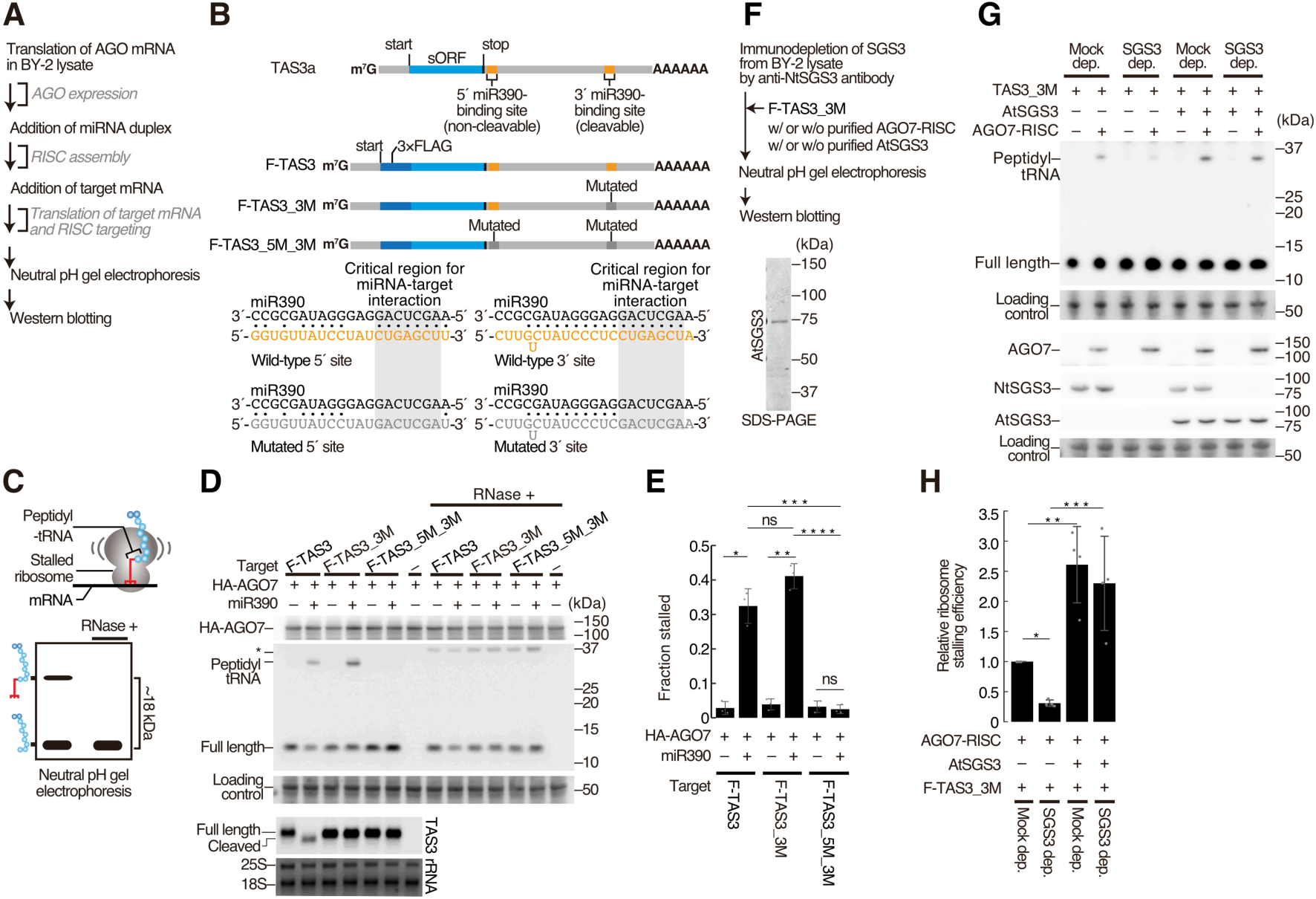
*In vitro* recapitulation of microRNA-mediated ribosome stalling. (A) Flowchart of the miRNA-mediated ribosome stalling assay *in vitro*. (B) (top) Schematic representation of TAS3a RNA and its 3×FLAG-tag fused variants. The orange and gray boxes indicate wild-type and mutated miR390-binding sites, respectively. (bottom) The base-pairing configurations between miR390 and the wild-type or mutated miR390-binding sites. The critical regions for the miRNA-target interaction are shown in the shaded boxes. (C) Schematic representation of SDS-PAGE in a neutral pH environment, to thus detect peptidyl-tRNAs. (D) Both AGO7-RISC and the 5′ binding site are required for ribosome stalling *in vitro*. After *in vitro* silencing assay, half of the reaction mixture was treated with RNase (RNase +), and used for PAGE followed by Western blotting. The full-length polypeptide and peptidyl-tRNA were detected by anti-FLAG antibody. 3×HA-AGO7 (HA-AGO7) was detected by anti-HA antibody. Total protein was stained using Ponceau S, and the ∼50 kDa bands were used as a loading control. The asterisk indicates the positions of the unexpected protein bands that appears with RNase treatment. (bottom) Northern blotting of TAS3 variants. Methylene blue-stained rRNA was used as a loading control. (E) Quantification of ribosome stalling efficiencies in (D). Fraction stalled was calculated using the following formula: Fraction stalled = peptidyl-tRNA/(full-length + peptidyl-tRNA). The mean values ± SD from three independent experiments are shown. Bonferroni-corrected P values from two-sided paired t-tests are as follows: *P = 0.03361; **P = 0.03817; ***P = 0.03809, ****P = 0.03174. (F) (top) Flowchart of the *in vitro* miRNA-mediated ribosome stalling assay with SGS3-immunodepleted lysate. (bottom) Coomassie brilliant blue staining of purified AtSGS3. (G) SGS3 promotes miRNA-mediated ribosome stalling *in vitro*. Endogenous NtSGS3, recombinant AtSGS3, and recombinant AGO7 were detected using anti-NtSGS3, anti-AtSGS3, and anti-AtAGO7 antibodies, respectively. See also Figure 2D legend. (H) Quantification of relative ribosome stalling efficiencies in (G). The signal intensity of peptidyl-tRNA/(full-length + peptidyl-tRNA) was normalized to the value of Mock dep. (AtSGS3 –). The mean values ± SD from four independent experiments are shown. Bonferroni-corrected P values from two-sided paired t-tests are as follows: *P = 0.00039; **P = 0.04433; ***P = 0.03889.

The two miR390-binding sites in TAS3a are functionally distinct; the 5′ possesses central mismatches that preclude RISC-mediated target cleavage but allow stable binding, whereas the 3′ miR390 binding site is centrally matched with the miR390 and thus cleaves the TAS3a RNA (Figure 2B) (Axtell et al., 2006). The adjacent 5′ binding site is essential for ribosome stalling. Mutations in the critical region for miRNA recognition of the 3′ miR390 binding site (Figure 2B, F-TAS3_3M) did not impair ribosome stalling, whereas also mutating the 5′ binding site (Figure 2B, F-TAS3_5M_3M) reduced stalling (Figure 2D and E). Thus, ribosome stalling requires base-pairing between miR390 in AGO7-RISC and the 5′ miR390-binding site in TAS3a. As the 3′ site mutation increased peptidyl-tRNA accumulation, presumably by stabilizing the mRNA since it is no longer cleaved (Figure 2D), we decided to use F-TAS3_3M for further experiments.

We next sought to investigate the impact of SGS3 on ribosome stalling. Because endogenous SGS3 (NtSGS3) is abundant in BY-2 cells (Yoshikawa et al., 2013), we immuno-depleted NtSGS3 from the lysate (Figure 2F) and found decreased ribosome stalling (Figure 2G and H). Supplementing with recombinant AtSGS3 markedly rescued ribosome stalling efficiency (Figure 2F–H), indicating that SGS3 is a critical and limiting factor for miRNA-mediated ribosome stalling. Taken altogether, our *in vitro* system faithfully recapitulated ribosome stalling triggered by AGO7-RISC and SGS3.

### SGS3 binding to the 3′ end of initiator microRNAs is required for ribosome pausing

The functional roles of AGO7-RISC and SGS3 prompted us to hypothesize that these two factors form a complex that promotes ribosome stalling. SGS3 is an RNA-binding protein that preferentially binds RNA duplexes with a 5′ overhang (Fukunaga and Doudna, 2009). In theory, such a substrate is formed between the 3′ end of miR390 within AGO7 and the 5′ end of the miR390-binding site. We therefore hypothesized that SGS3 directly interacts with the end of the dsRNA protruding from AGO7. To test this scenario, we first examined the interaction between SGS3 and AGO7-RISC. The FLAG-tagged AGO7 mRNA was translated in the BY-2 cell lysate, then the miR390 duplex was added to program RISC. After further incubation with TAS3 mRNAs, the reaction mixture was used for co-immunoprecipitation with anti-FLAG antibody (Figure 3A). This assay revealed that endogenous NtSGS3 binds AGO7-RISC only in the presence of both miR390 and TAS3 variants with a wild-type 5′ site (Figure S4A). Remarkably, introducing mismatches at the 5′ end of the miR390-binding site (Figure 3B, TAS3_5endM_3M) or using a miR390 variant that is one-nucleotide shorter (20 nt) (Figure 3B), which is not predicted to protrude from AGO7, disrupted the interaction between NtSGS3 and AGO7-RISC (Figure 3B–D, Figure S4B). These results strongly support a model where SGS3 forms a complex with AGO7-RISC via dsRNA with a 5′ overhang formed at the 3′ end of miR390.

**Figure 3.**
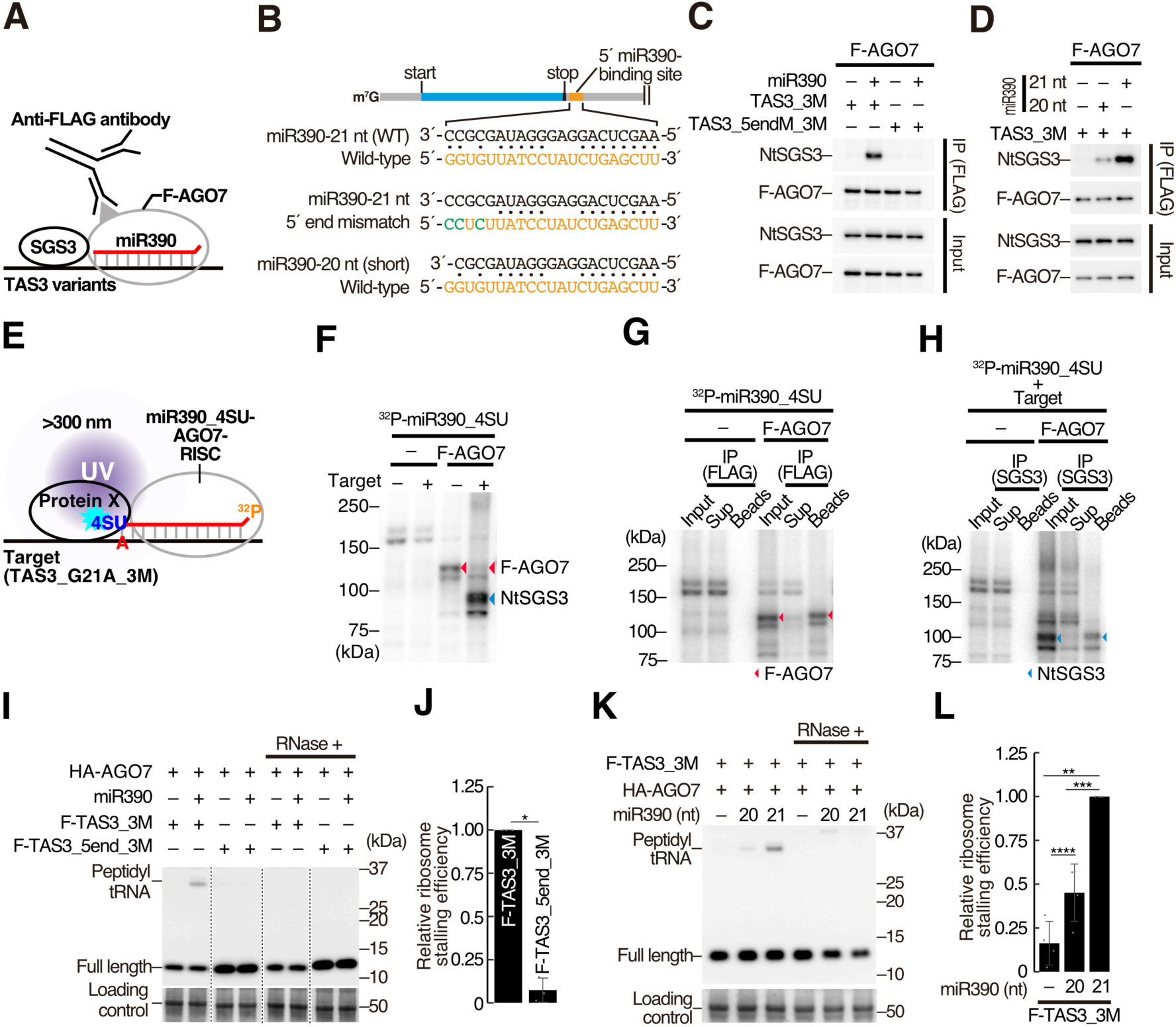
SGS3 binding to the 3′ end of miR390 is required for the ribosome pausing. (A) Schematic representation of co-immunoprecipitation assay with anti-FLAG antibody in the presence of F-AGO7, miR390, and TAS3 variants. (B) Base-pairing configurations. (top) miR390 and wild-type 5′ miR390-binding site. (middle) miR390 and 5′ site with 5′ end mismatches (5endM). (bottom) 20-nt miR390 and the wild-type 5′ site. The mutated nucleotides are shown in green. (C) 5′ end mismatches in the 5′ site disrupt interaction between SGS3 and AGO7. (D) The use of a short miR390 variant (20 nt) disrupted interaction between SGS3 and AGO7. (E) An overview of the UV crosslink experiment. AGO7 was programmed with the 5′-radiolabeled miR390 variant bearing the 3′ 4-thio-U (^32^P-miR390_4SU) in BY-2 lysate, and further incubated with the TAS3 variant (TAS3_G21A_3M). The reaction mixture was analyzed using 10% SDS-PAGE and crosslinked proteins were detected using phosphorimaging. The ellipse indicates neighboring proteins. (F) miR390-loaded AGO7 directly interacts with NtSGS3 in the presence of TAS3_G21A_3M. 5′ end- radiolabeled miR390 with a 3′ 4-thio-U was incubated in BY-2 lysate in the presence or absence of F-AGO7 and target RNA (TAS3_G21A_3M), crosslinked by UV light (>300 nm), then analyzed by SDS-PAGE. The red and blue arrowheads indicate AGO7 and NtSGS3, respectively (See also g and h). (G) Detection of F-AGO7 by UV crosslinking. The 5′ end-radiolabeled miR390 with a 3′ 4-thio-U was incubated in BY-2 lysate in the presence of F-AGO7, crosslinked by UV light (>300 nm), immunoprecipitated using anti-FLAG antibody, and then analyzed by SDS-PAGE. F-AGO7 was efficiently crosslinked to 4-thio-U at the 3′ end of miR390. (H) Detection of NtSGS3 by UV crosslinking. 5′ end-radiolabeled miR390 with a 3′ 4-thio-U was incubated in BY-2 lysate in the presence of F-AGO7 and target RNA (TAS3_G21A_3M), crosslinked by UV light (>300 nm), immunoprecipitated by anti-NtSGS3 antibody, and then analyzed by SDS-PAGE. NtSGS3 was efficiently crosslinked to 4-thio-U at the 3′ end of miR390 in the presence of the target RNA. (I) and (K) *in vitro* ribosome stalling experiments. Mismatches at the 5′ end of miR390-binding site or the use of 20-nt miR390 decreased stalled ribosomes. See also the legend of Figure 2D. (J) and (L) Quantification of relative ribosome stalling efficiencies in (I) and (K), respectively. The signal intensity of peptidyl-tRNA/(full-length + peptidyl-tRNA) was normalized to the value of F-TAS3_3M (I) or miR390 (21 nt) (K). The mean values ± SD from three (J) and four (L) independent experiments are shown, respectively. P value from two-sided paired t-tests are as follows: *P = 0.00190 (J). Bonferroni-corrected P values from two-sided paired t-tests are as follows: **P = 0.00270; ***P = 0.02034; ****P = 0.04343 (L).

To test whether SGS3 directly interacts with the 3′ end of miR390 on the *TAS3* RNA, we performed a site-specific UV crosslinking assay, in which molecules neighboring the 3′ end of miR390 can be captured. We first substituted the 3′ end cytidine of miR390 with a photo-reactive 4-thiouridine (Figure S4C, miR390_4SU), and restored base-pairing using a TAS3a variant with a G-to-A substitution at the 5′ miR390 binding site (Figure S4C, G21A substitution). This variant successfully rescued the interaction between miR390_21_4SU-loaded AGO7 and NtSGS3 (Figure S4C and D). In this context of the reporter, proteins crosslinked to 5′ radiolabeled miR390_21_4SU were separated on an SDS-PAGE gel (Figure 3E). In the absence of the target RNA, a specific band appeared at around 120 kDa (Figure 3F, red arrowhead). Immunoprecipitation using the anti-FLAG antibody revealed that the band corresponds to F-AGO7 (Figure 3G, red arrowheads). Strikingly, addition of the target RNA changed the crosslinked protein from AGO7 to a ∼90 kDa protein (Figure 3F). This protein was immunoprecipitated with anti- NtSGS3 antibody (Figure 3H, Beads), and specifically depleted from the supernatant of the lysate after immunoprecipitation (Figure 3H, Sup), corroborating the identity of this ∼90 kDa crosslinked protein as NtSGS3. These results indicate that target binding alters protein interactions at the 3′ end of miR390, switching them from AGO7 to SGS3, likely via conformational changes in AGO7-RISC.

To test if the physical interaction between SGS3 and AGO7-RISC is critical for ribosome pausing, we performed *in vitro* ribosome stalling experiments under conditions where SGS3 fails to bind AGO7-RISC using reporter variant F-TAS3_5endM_3M or the short 20-nt version of miR390. In both cases, stalling efficiencies were significantly decreased (Figure 3I–L). Taken together, we find a direct interaction between SGS3 and the 3′ end of 21-nt miR390 bound to AGO7-RISC is necessary for ribosome stalling on TAS3 mRNA.

It is worth noting that the required length of miRNA for SGS3 binding and ribosome pausing may differ between partner AGO proteins. In contrast to AGO7, AGO1—bound by most miRNAs—requires a 22-nt long miR173 for both SGS3 interaction and ribosome stalling (Figure 4A–D) (Yoshikawa et al., 2013). As miRNAs are typically 21-nt long, plants may have evolved a AGO1 structure that fully encapsulates the 21-nt miRNAs, thus limiting promiscuous SGS3 binding and ribosome stalling (Figure 4C and D). Importantly, we find that ribosome pausing occurs even if TAS1 is cleaved by AGO1-RISC loaded with 22-nt miR173 (Figure 4D and E). This is not limited to the TAS1 and miR173-AGO1 pair. AGO7-miR390-SGS3 complex also stalls ribosomes on the cleavable binding site which has perfect complementarity to miR390 (Figure S5A–C). Because AGO-miRNA-SGS3 complex holds and stabilizes both 5′ and 3′ RNA fragments after target cleavage (Figure 4E) (Yoshikawa et al., 2013), we reasoned that SGS3 and RISC can stay on the cleaved targets long enough to stall ribosomes.

**Figure 4.**
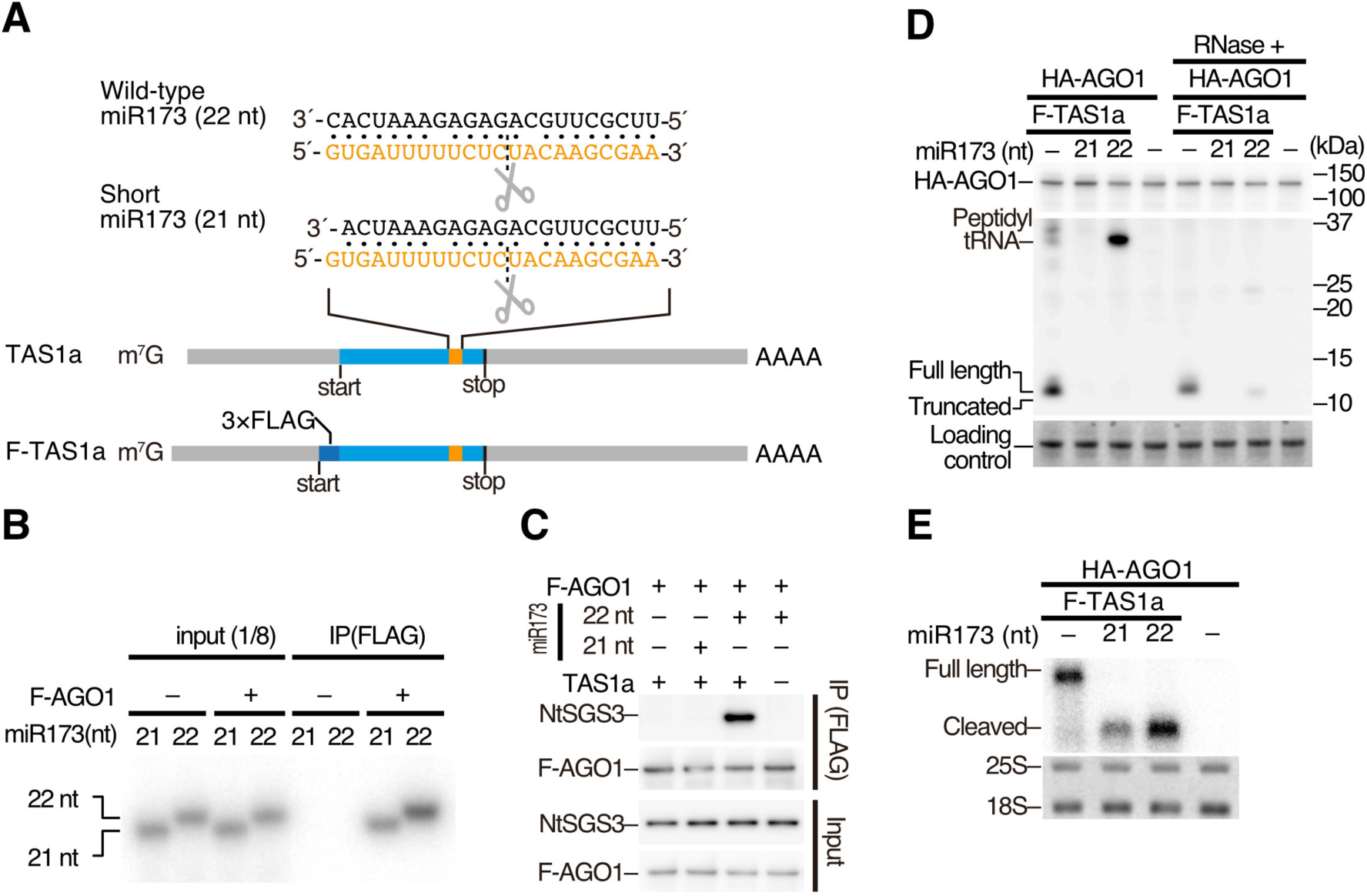
AGO1 loaded with 22-nt miR173 efficiently stalls ribosome. (A) (top) Base-pairing configurations between 22/21-nt miR173 and the miR173-binding site in TAS1a. (bottom) Schematic representation of TAS1a RNA and its 3×FLAG-tag fused variant. (B) *In vitro* RISC assembly with AGO1 and radiolabeled 21 and 22-nt miR173 duplexes. After translation of 3×FLAG-AGO1 (F-AGO1) mRNA *in vitro*, the radiolabeled miR173 duplex was added and further incubated for RISC assembly. Then, F-AGO1 was immunoprecipitated with anti-FLAG antibody. The co-immunoprecipitated miR173 was analyzed by denaturing PAGE. Both 21- and 22-nt miR173 duplexes were incorporated into AGO1. (C) Co-immunoprecipitation experiments with 3×FLAG-AGO1 in the presence of 21 or 22-nt miR173 duplex and TAS1a RNA. AGO1-RISC loaded with 22- nt miR173 interacts with NtSGS3 in the presence of TAS1a RNA. In contrast, 21-nt miR173 failed to promote the interaction between AGO1 and NtSGS3. (D) *In vitro* ribosome stalling experiments. Peptidyl-tRNA was accumulated in the presence of AGO1-RISC loaded with 22-nt miR173, while no peptidyl-tRNA was observed in the presence of that with 21-nt miR173. (E) Northern blotting of TAS1 reporter RNAs. TAS1 was efficiently cleaved by AGO1-RISC loaded with 21- and 22-nt miR173. Methylene blue-stained rRNA was used as a loading control.

### Ribosome stalling is not an essential event but a positive modulator for the production of secondary siRNAs

The striking correspondence between ribosome stalling and TAS precursors (Figure 1 and S1B–D) led us to hypothesize that ribosome pausing by the SGS3-miRNA complex promotes tasiRNA production. So far, several studies have focused on the relationship between translation and tasiRNA biogenesis (Zhang et al., 2012; Hou et al., 2016; Li et al., 2016; Yoshikawa et al., 2016; Bazin et al., 2017). However, it is still controversial if positioning of the miRNA-binding site in the CDS or near the stop codon is important for tasiRNA production (Zhang et al., 2012; Yoshikawa et al., 2016; Bazin et al., 2017). For example, previous quantitative RT-PCR (qRT-PCR) experiments showed no significant changes in tasiRNA production between the wild-type TAS3 and a mutant TAS3 that possesses an early stop codon located far upstream of the 5′ miR390 binding site (Bazin et al., 2017), suggesting that ribosome stalling has no impact on the tasiRNA biogenesis. To carefully assess the impact of ribosome stalling on tasiRNA biogenesis, we first attempted to construct TAS3 variants with no ribosome stalling that retain binding to AGO7 and SGS3. Such variants were obtained by inserting 4 or more nucleotides between the stop codon and 5′ miR390 binding site in TAS3 (Figure 5A and B, S6). As ribosomes stall one-codon upstream of the stop codon in TAS3, we reasoned that these insertions promote normal translation termination without interfering in the binding between AGO7-RISC and SGS3.

**Figure 5.**
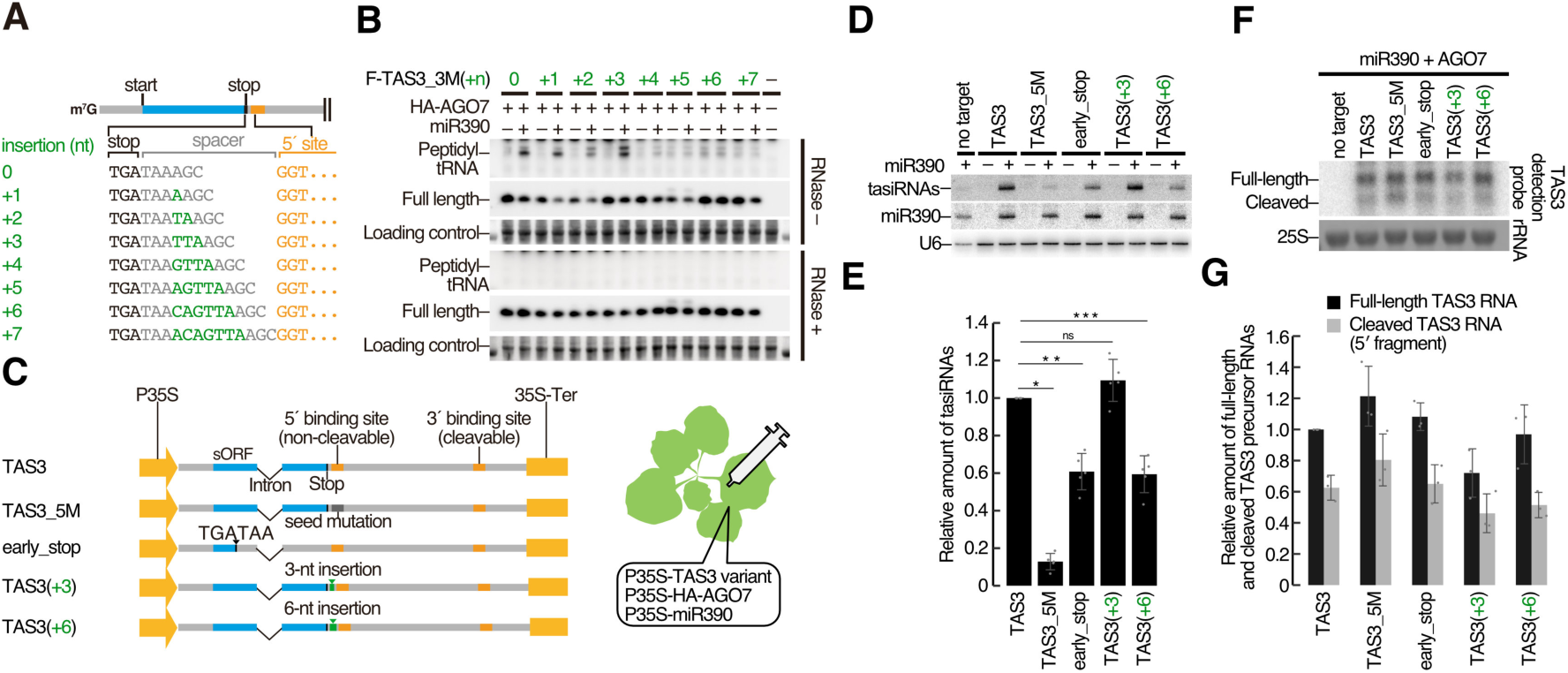
Ribosome stalling is not an essential event but an enhancer for the production of secondary siRNAs. (A) Schematic representation of TAS3 variants with different nucleotide insertions (green) between the stop codon (black) and the 5′ miR390-binding site (orange). (B) *In vitro* ribosome stalling experiments. Insertions of over 3 nucleotides decreased stalled ribosomes. After *in vitro* silencing assay, half of the reaction mixture was treated with RNase (RNase +), and used for PAGE followed by Western blotting. The full-length polypeptide and peptidyl-tRNA were detected by anti-FLAG antibody. Total protein was stained using Ponceau S, and the ∼50 kDa bands were used as a loading control. (C) Schematic representation of plasmids carrying TAS3 variants used in the *Nicotiana benthamiana (N. benthamiana)* transient assay. P35S and 35S-Ter indicate Cauliflower mosaic virus (CaMV) 35S promoter and terminator, respectively. Leaves of *N. benthamiana* plants were infiltrated with a mixture of *Agrobacterium tumefaciens* cultures harboring P35S-HA-AGO7, P35S-TAS3 variants, and P35S-miR390 or empty vector. Leaves were harvested at 2-day post infiltration and used for Northern blotting to detect secondary siRNAs and the sense strand of TAS3 mRNAs. (D) Northern blotting of secondary siRNAs from TAS3 and its variants, miR390, and U6 RNAs. (E) Quantification of the secondary siRNAs in (D). The signal intensity of tasiRNAs was calibrated with miR390, and normalized to the value of TAS3 (miR390 +). The mean values ± SD from five independent experiments are shown. Bonferroni-corrected P values from two-sided paired t-tests are as follows: *P = 5.96313E-06; **P = 0.00332; ***P = 0.00329. A positive correlation was observed between tasiRNA biogenesis and miR390-mediated ribosome stalling. (F) Northern blotting of the full-length TAS3 RNAs and the 5′ cleaved fragments. Methylene blue-stained rRNA was used as a loading control. (G) The signal intensity of TAS3 RNA/rRNA was normalized to the value of full-length TAS3. The mean values ± SD from three independent experiments are shown. No correlation was observed between ribosome stalling and accumulation of the sense strand of TAS3 RNAs.

To test the hypothesis that ribosome pausing promotes the production of tasiRNAs, we compared tasiRNA accumulation in different TAS3 variants in *Nicotiana benthamiana* leaves. We opted to use Northern blotting for accurate detection of the secondary siRNAs, because this method can distinguish the canonical secondary siRNAs from the non-specific RNA fragments derived from the TAS3 reporters by size. Co-expression of miR390 and AGO7 efficiently produced 21-nt tasiRNAs, compared with the 5′ miR390 binding site mutant (TAS3_5M) (Figure 5C–E). Thus, our transient assay successfully recapitulated canonical TAS3 tasiRNA biogenesis. Importantly, placing the 5′ miR390 binding site 6-nucleotide away (Figure 5C, TAS3+6) significantly reduced tasiRNA production to ∼60% (Figure 5D and E), suggesting that clearance of stalled ribosomes impairs efficient tasiRNA production. In contrast, tasiRNA production from a TAS3 variant with a 3-nucleotide insertion (Figure 5C, TAS+3), which still stalls ribosomes, was comparable to that from wild-type TAS3 (Figure 5D and E). To confirm if ribosome stalling enhances the tasiRNA biogenesis, we introduced artificial tandem stop codons at the ∼120 nt upstream of the miR390 target site (early_stop), which forces ribosomes to terminate without stalling (Figure 5C). In contrast to the previous report (Bazin et al., 2017), our quantitative Northern blotting revealed that the tandem early stop codons significantly reduced tasiRNA production to ∼60%, similarly to TAS3+6 (Figure 5D and E). These data were not explained by changes in precursor *TAS3* abundance (Figure 5F and G). Altogether, we conclude that ribosome stalling regulates secondary siRNA production in a manner different from stabilization of mRNAs.

## Discussion

Here, we find that the dsRNA-binding protein SGS3 forms ribosome stalling complexes on the protruding end of the dsRNA formed between the TAS RNAs and miR390-AGO7 or 22-nt miR173-AGO1-RISC (Figure 6). In general, the ribosome displaces RNA binding proteins bound to mRNAs during elongation (Halstead et al., 2016), suggesting that SGS3 imposes an extreme barrier for trailing ribosomes. A recent study suggested that unconventional base-pairing between human miRNAs and target sites cause transient ribosome stalling (Zhang et al., 2018). Although the precise stalling mechanism remains unclear in animals, there may be an RNA-binding protein(s) that protects the 3′ end of miRNA from the helicase activity of ribosomes.

**Figure 6.**
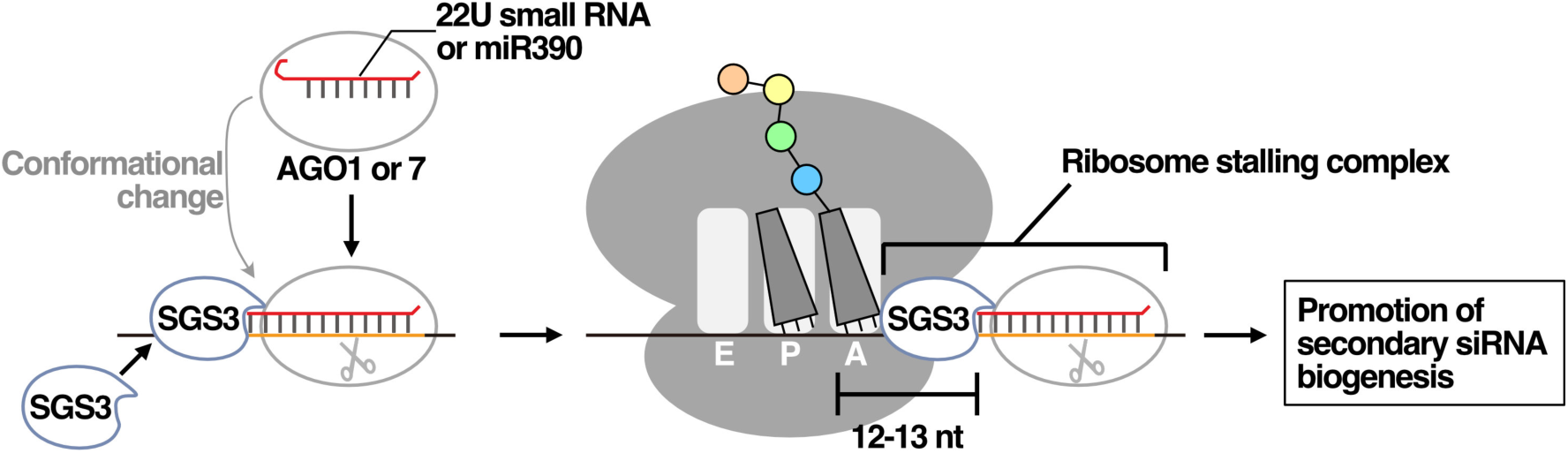
A model for ribosome stalling caused by SGS3-miRNA-Argonaute complex and its role in secondary siRNA biogenesis. Target binding causes dynamic conformational changes in 22U-AGO1-RISC or miR390-AGO7-RISC, resulting in protrusion of the 3′ end of the small RNA from the RISC complex. SGS3 directly binds the dsRNA formed between the 3′ side of the small RNA and the 5′ side of the target site. The SGS3- small RNA complex stalls ribosomes at 12–13 nt upstream of the binding sites. This ribosome stalling stimulates secondary siRNA production in a manner different from mRNA stabilization.

The accumulating evidence suggests that translation alters the biogenesis of TAS3 tasiRNAs (Li et al., 2016; Bazin et al., 2017). A previous study showed that the position of the start site is critical for the stability of TAS3 mRNAs and tasiRNA biogenesis (Bazin et al., 2017). We here demonstrate that ribosome stalling enhances TAS3 tasiRNA biogenesis (Figure 5D, E and 6). This is supported from an evolutionary standpoint; many plant species have the 5′ miR390 binding site just downstream the stop codon or in the CDS of TAS3 (Table S2). This positive effect of ribosome pausing for secondary siRNA production may not be limited to TAS3 genes. It was previously shown that a miR173 binding site located within ORF also enhances secondary siRNA biogenesis (Zhang et al., 2012; Yoshikawa et al., 2016), suggesting that ribosome stalling promotes tasiRNA biogenesis on TAS1/2 genes and other 22-nt miRNA target genes. On the other hand, ribosome stalling is not a prerequisite for triggering secondary siRNA biogenesis. Indeed, TAS3+6 and early_stop still produced tasiRNAs in our transient assays with *Nicotiana benthamiana* plants (Figure 5D and E). Thus, although stalled ribosomes positively regulate tasiRNA production, SGS3-miRNA-AGO complex can trigger tasiRNAs independently of translational arrest (Figure 6). The molecular details of how ribosome stalling enhances tasiRNA production warrant future studies. Given that arrest peptide-mediated ribosome-pausing induces changes in mRNA localization in animal cells (Yanagitani et al., 2011), we suggest that miRNA-mediated ribosome pausing may facilitate the delivery of the tasiRNA precursors to a secondary siRNA “factory”, such as the siRNA body (Jouannet et al., 2012).

We observed SGS3- and RISC-dependent ribosome stalling in five TAS loci in *Arabidopsis* (Figure 1). However, they may be just a tip of the iceberg of miRNA-mediated ribosome pausing. There are many DNA regions named PHAS loci that produce phased secondary siRNAs (phasiRNAs) by the same mechanism as TAS loci (Liu et al., 2020). Although our ribosome profiling failed to detect obvious ribosome stalling 11–14 nt upstream of miRNA binding sites in known PHAS loci (Figure 1 and Table S1), more sensitive methods like single-molecule imaging (Ruijtenberg et al., 2020) may reveal ribosome stalling on the miRNA-bound targets. In addition to miRNAs, siRNAs may also induce ribosome stalling. A recent study demonstrates that 22-nt siRNAs, which have the potential to recruit SGS3, accumulate upon environmental stress, trigger the RNA silencing amplification, and mediate translational repression (Wu et al., 2020). Such 22- nt siRNAs are also induced by viral infection (Mourrain et al., 2000; Akbergenov et al., 2006; Deleris et al., 2006; Diaz-Pendon et al., 2007; Garcia-Ruiz et al., 2010). Therefore, SGS3- and miRNA/siRNA-mediated ribosome stalling is likely to have an impact on a wider range of cellular processes such as stress adaptation and antiviral immunity in plants.

## Methods

### General methods

Preparation of tobacco BY-2 lysate, substrate mixture (containing ATP, ATP-regeneration system, and amino acid mixture), 1×lysis buffer [30 mM HEPES-KOH (pH 7.4), 100 mM potassium acetate, 2 mM magnesium acetate], and microRNA duplexes (Table S3) have been previously described in detail (Tomari and Iwakawa, 2017). mRNAs were transcribed *in vitro* from NotI- (for plasmids with the prefix “pBYL-”) or XhoI- (for plasmids with the prefix “pUC57-”) digested plasmids or PCR products using the AmpliScribe T7 High Yield Transcription Kit (Lucigen), followed by capping with ScriptCap m^7^G Capping System (Cell Script). Poly(A)-tails were added to transcripts from pUC57-plasmids or PCR products using the T7 promoter by A-Plus Poly(A) Polymerase Tailing Kit (Cell Script). Anti-AtSGS3 (diluted at 1:3000) and anti-AtAGO7 antibodies (diluted at 1:3000) were raised in rabbits using synthetic peptides (NH_2_- MSSRAGPMSKEKNVQGGC-COOH) and (NH_2_-IPSSKSRTPLLHKPYHHC-COOH) as antigens respectively, and affinity-purified (Medical & Biological Laboratories).

### Plants and growth conditions

*Arabidopsis thaliana* wild-type (Col-0) and the *sgs3-11* mutant (Peragine et al., 2004) were used in this study. Seeds were incubated in 70% EtOH at room temperature for 2 min, sterilized with liquid sodium hypochlorite, washed 5 times in sterile water, sown on filter paper (Whatman No.2), laid on Murashige and Skoog (MS)-agar plates (1×MS salt, 1% sucrose, 1% agar, pH 5.7) and incubated at 4°C for 3 days. After vernalization, the plates were incubated at 22°C for 3 days under continuous LED light (LC-LED450W, TAITEC).

### Ribosome profiling

Briefly, 0.2 g of frozen seedlings and 400 µl of *Arabidopsis* lysis buffer (100 mM Tris-HCl pH 7.5, 40 mM KCl, 20 mM MgCl_2_, 1 mM DTT, 100 µg/ml cycloheximide and 1% Triton X-100) were crushed into a powder using the Multi-beads shocker (Yasui Kikai). The 3000 × g supernatant of the lysate was mixed with 25 µl of Turbo DNase (Thermo Fisher Scientific) and incubated on ice for 10 min. RNA concentration was measured with a Qubit RNA BR Assay Kit (Thermo Fisher Scientific). Ribosome footprints ranging between 17 and 34 nt were gel-purified and subsequent library preparation were executed as previously described (McGlincy and Ingolia, 2017; Kurihara et al., 2018). Two libraries from two biological replicates (WT_rep1, WT_rep2, sgs3_rep1 and sgs3_rep2) were sequenced on a HiSeq4000 (Illumina). 24 to 29 nt footprints were mapped onto the TAIR10 *Arabidopsis thaliana* genome sequence, excluding rRNA/tRNAs. Empirically, A-site position was estimated as 11 for 24 nt, 12 for 25 nt, 13 for 26 nt, 14 for 27 nt, 15 for 28 nt, 16 for 29 nt, based on the homogeneous 5′ end of the reads. The relative ribosome occupancy *r* at position *j* in an ORF of gene *g* of length *l* is defined as follows:

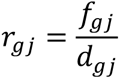

where

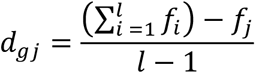

*f_gj_* is the footprint at position *j* in a ORF of gene *g*. *r_gj_* is a ratio of *f_gj_* to the average footprint across nucleotide positions on the ORF of the same gene, *d_gj_*.

### microRNA target prediction

The targets of mature *Arabidopsis* microRNA sequences [miRbase (miRbase20) (Kozomara and Griffiths-Jones, 2011; Kozomara and Griffiths-Jones, 2014)] were predicted using the psRNATarget server (Dai and Zhao, 2011; Dai et al., 2018) with the following settings: # of top targets = 15, Expectation = 3, Seed region = 2-8 nt.

### RNA-seq

Total RNA was extracted from seedlings with Trizol (Thermo Fisher Scientific). Library construction and deep sequencing were performed by AnnoRoad in Beijing. Reads were mapped to the transcripts of *Arabidopsis thaliana* (derived from TAIR10, ver. 10 released on 2010 in psRNATarget server (Dai and Zhao, 2011; Dai et al., 2018)) by Bowtie2(Langmead and Salzberg, 2012). Sam files were converted to bam files using SAMtools (Li et al., 2009) and then to bed files with BEDTools (Quinlan and Hall, 2010). BEDtools (Quinlan and Hall, 2010) was used to calculate the depth of coverage for every base across mRNAs shown in Figure 1B, C, S1B–D.

### Plasmid construction

The following constructs used in this study have been previously described: pBYL2 (Mine et al., 2010), pBYL-AGO1 (Endo et al., 2013), pBYL-AGO7 (Endo et al., 2013), pBYL-3×FLAG-AGO7 (Endo et al., 2013), pBYL-3×FLAG-AGO1 (Endo et al., 2013), pBYL-3×FLAG-SUMO-AtAGO1 (Iwakawa and Tomari, 2013), pAT006 (Tsuzuki et al., 2014), pMDC32 (Curtis and Grossniklaus, 2003), pMDC-Tas3a (Montgomery et al., 2008), pMDC-HA-AGO7 (Montgomery et al., 2008), pMDC-miR390 (Montgomery et al., 2008). The DNA fragments used for plasmid construction are listed in Table S4.

#### pBYL-3×HA

A DNA fragment containing the T7 promoter, 5′ UTR of *Arabidopsis thaliana* alcohol dehydrogenase 1 and 3×HA tag (T7_ADH_5UTR_*3×HA,* Table S4) was cloned into XbaI/AscI-digested pBYL2 vector using the HiFi DNA Assembly Cloning kit (New England Biolabs).

#### pBYL-3×HA-AGO7

A DNA fragment containing AGO7 ORF was amplified by PCR with pBYL-AGO7(Endo et al., 2013) using primers oligoE1 and oligoE2, digested by AscI, and cloned into AscI-digested pBYL-3×HA vector by ligation.

#### pBYL-3×HA-AGO1

A PCR fragment with AGO1 ORF following 3×HA tag was amplified by overlap extension PCR with pBYL-AGO1 (Endo et al., 2013) as template using primers oligo1118 and oligo1094. The fragment was cloned into AscI-digested pBYL2 vector via HiFi DNA Assembly Cloning kit (New England Biolabs).

#### pUC57-TAS3

The TAS3a sequence (AT3G17185.1) following T7 promoter (T7_TAS3a, Table S4) was inserted into EcoRV-digested pUC57 vector via GenScript gene synthesis service.

#### pUC57-F-TAS3

Three DNA fragments were prepared by PCR: TAS3a_5′ UTR fragment amplified from pUC57-TAS3 using primers oligo1062 and oligo1063, 3×FLAG tag sequence amplified using two oligos, oligo1064 and oligo512 and the TAS3a ORF amplified from pUC57- TAS3 using primers, oligo1065 and oligo1066. The three DNA fragments were cloned into SacII/XhoI-digested pUC57-TAS3a via HiFi DNA Assembly Cloning kit (New England Biolabs).

#### pUC57-F-TAS3_3M

Seven nucleotide mismatches were introduced into the 3′ miR390 binding site (Figure 2C) in pUC57-F-TAS3 by site directed mutagenesis using primers oligo1073 and oligo1074.

#### pUC57-F-TAS3_5M_3M

Seven nucleotide mismatches were introduced into the 5′ miR390 binding site (Figure 2C) in pUC57-F-TAS3_3M by site directed mutagenesis using primers oligo 1099 and oligo1100.

#### pUC57-F-TAS3_3M(+1), pUC57-F-TAS3_3M(+2), pUC57-F-TAS3_3M(+3), pUC57-F-TAS3_3M(+4), pUC57-F-TAS3_3M(+5), pUC57-F-TAS3_3M(+6) and pUC57-F-TAS3_3M(+7)

One to six nucleotides, as shown in Figure 5A, were inserted between the stop codon of the short ORF and 5′ miR390 binding site in pUC57-F-TAS3_3M by site directed PCR using primer pairs of oligo1180-oligo1181, oligo1182-oligo1183, oligo1161-oligo1162, oligo1163-oligo1164, oligo1165-oligo1166, oligo1167-oligo1168 and oligo1169-oligo1170, respectively.

#### pEU-6×His-SBP-SUMO-AtSGS3

Two DNA fragments were prepared by PCR: 6×His-SBP-SUMOstar-tag fragment amplified from pASW-SUMO-AtRDR6 (Opt) (Baeg et al., 2017) using oligo1044 and oligo1039 and SGS ORF fragment amplified from cDNA of *Arabidopsis thaliana* using oligoK1 and oligoK2. The two DNA fragments were inserted into EcoRV/SmaI-digested pEU-E01-MCS vector via HiFi DNA Assembly Cloning kit (New England Biolabs).

#### pBYL-3×FLAG-SUMOstar-tag-AGO7

Two PCR products were prepared by PCR: 3×FLAG-SUMOstar-tag fragment amplified from pBYL-3×FLAG-SUMO-AtAGO1 (Iwakawa and Tomari, 2013) using primers oligo955 and oligo1039 and AGO7 fragment amplified from pBYL-AGO7 (Endo et al., 2013) using primers oligo1159 and oligo1160. The two fragments were cloned into AscI-digested pBYL2 vector (Mine et al., 2010) via HiFi DNA Assembly Cloning kit (New England Biolabs).

#### pUC57-F-TAS3_5endM_3M and pUC57-F-TAS3_5P_3M

The 5′ miR390-binding site in pUC57-F-TAS3_3M was replaced by the sequences shown in Figure 3B and Figure S5 by site directed mutagenesis using primer pairs oligo1101- oligo1102 and oligo 1106-oligo1107, respectively.

#### pUC57-TAS3_3M, pUC57-TAS3_5endM_3M, pUC57-TAS3_5P_3M, pUC57-TAS3_M(+1), pUC57-TAS3_M(+2), pUC57-TAS3_M(+3), pUC57-TAS3_M(+4), pUC57-TAS3_M(+5), pUC57-TAS3_M(+6) and pUC57-TAS3_M(+7)

The 3×FLAG tag sequences were removed from the corresponding pUC57-F-TAS3 constructs shown above by site directed mutagenesis using primers oligo1197 and oligo1198.

#### pUC57-TAS3_G21A_3M

The 5′ terminal G nucleotide of 5′ miR390-binding site in pUC57-TAS3_3M was substituted to A by site directed mutagenesis using primers oligo1220 and oligo1221.

#### pCR-Blunt II-TOPO_TAS1a

TAS1a PCR product was amplified from cDNA corresponding to *Arabidopsis* seedling total RNA using oligoA1 and oligoA2 for the TAS1a sequence and cloned into pCR Blunt II-TOPO vector (Invitrogen, #45-0245).

#### pCR-Blunt II-TOPO_3×FLAG-TAS1a

Three PCR fragments were prepared from pCR-Blunt II-TOPO_TAS1a: TOPO-TAS1a 5′ UTR fragment amplified with oligoA3 and oligoA4, FLAG-TAS1a fragment amplified with oligoA5 and oligoA6 and ORF-3′ UTR-TOPO fragment amplified with oligoA7 and oligoA8. To insert the 3×FLAG sequence directly in front of ORF1, the above three PCR fragments were cloned into XhoI/SpeI-digested pCR Blunt II-TOPO vector (Invitrogen) using the HiFi DNA Assembly Cloning kit (New England Biolabs).

##### T7-TAS1a and T7-F-Tas1a

T7-TAS1a and T7-F-Tas1a DNA templates were amplified from pCR-Blunt II- TOPO_TAS1a and pCR Blunt II-TOPO-3xFLAG-TAS1a, respectively, using a forward primer containing T7 polymerase binding site (oligoA9) and a reverse primer with poly(A) tail (oligoA10).

#### pAT006-TAS3a-PDS_ full-length

A TAS3a fragments with a full-length 5′ UTR, a natural intron and tandem synthetic-tasiRNAs in the 5′ D7[+] and 5′ D8[+] positions (TAS3aPDS2) was synthesized via GeneArt Strings DNA Fragments service (invitrogen), gel-purified and cloned into SalI/SpeI-digested pAT006 (Tsuzuki et al., 2014) vector via HiFi DNAAssembly Cloning kit (New England Biolabs).

#### pMDC32_TAS3

Two PCR products were amplified: fragment A from pAT006-TAS3a-PDS_full-length using primers oligo1201 and oligo1202 and fragment B from pMDC-Tas3a (Montgomery et al., 2008) using primers oligo1203 and oligo1204. The two fragments were cloned into KpnI/SpeI-digested pMDC32 vector via HiFi DNA Assembly Cloning kit (New England Biolabs).

#### pMDC32_TAS3_5M

Two PCR products were amplified: fragment A from pAT006-TAS3a-PDS_full-length using primers oligo1201 and oligo1209 and fragment B from pMDC-Tas3a (Montgomery et al., 2008) using primers oligo1210 and oligo1204. The two fragments were cloned into KpnI/SpeI-digested pMDC32 vector via HiFi DNA Assembly Cloning kit (New England Biolabs).

#### pMDC32_early_stop

Two PCR products were amplified: fragment A from pAT006-TAS3a-PDS_full-length using primers oligo1201 and oligo1211 and fragment B from pMDC-Tas3a (Montgomery et al., 2008) using primers oligo1212 and oligo1204. The two fragments were cloned into KpnI/SpeI-digested pMDC32 vector via HiFi DNA Assembly Cloning kit (New England Biolabs).

#### pMDC32_TAS3(+3)

Two PCR products were amplified: fragment A from pAT006-TAS3a-PDS_full-length using primers oligo1201 and oligo1207 and fragment B from pMDC-Tas3a (Montgomery et al., 2008) using primers oligo1208 and oligo1204. The two fragments were cloned into KpnI/SpeI-digested pMDC32 vector via HiFi DNA Assembly Cloning kit (New England Biolabs).

#### pMDC32_TAS3(+6)

Two PCR products were amplified: fragment A from pAT006-TAS3a-PDS_full-length using primers oligo1201 and oligo1205 and fragment B from pMDC-Tas3a (Montgomery et al., 2008) using primers oligo1206 and oligo1204. The two fragments were cloned into KpnI/SpeI-digested pMDC32 vector via HiFi DNA Assembly Cloning kit (New England Biolabs).

### Production of recombinant AtSGS3 protein

Recombinant AtSGS3 proteins were expressed using the Premium PLUS Expression kit (Cell-Free Sciences) with pEU-6×His-SBP-SUMO-AtSGS3 according to manufacturer instructions. The protein was affinity purified with streptavidin sepharose high performance beads (GE Healthcare), washed three times with l × lysis buffer containing 200 mM NaCl and 0.1% TritonX-100, rinsed once with l × lysis buffer containing 20% glycerol and 1mM DTT and eluted by l × lysis buffer containing 20% glycerol, 1 mM DTT and 0.05 U/µl of SUMOstar protease. Protein concentration was determined using SDS-PAGE with defined dilutions of BSA as concentration standards.

### *In vitro* RNA silencing assay, NuPAGE and Western blotting

Typically, 7.5 µl of BY-2 lysate, 3.75 µl of substrate mixture, and 0.75 µl of 300 nM AGO mRNAs were mixed and incubated at 25°C for 30 min. To assemble RISC, 1.5 µl of 1.5 µM miR390 or miR173 duplex was added to the reaction mixture and incubated at 25°C for 90 min. Then, 1.5 µl of 100 nM TAS3a or TAS1a variant was added and further incubated at 25°C for 10–60 min. For RNase treatment, 5 µl of the reaction was treated with 1 µl of RNase mixture (10% RNase A, Sigma + 20% RNase One, Promega), incubated at 37°C for 10 min and then mixed with 6 µl of 2 × SDS-PAGE buffer. For the control, 1 µl sterile water was used instead of RNase mixture. The samples were run on NuPAGE Bis-Tris Precast Gel (Thermo Fisher Scientific) at 200 V for ∼30 min in 1 × NuPAGE MES SDS Buffer (Thermo Fisher Scientific) and transferred onto PVDF membrane. Western blotting was performed as previously described (Tomari and Iwakawa, 2017) with modifications. The membrane was blocked in TBST containing 1.0% nonfat dried milk (w/v) for 30 min. Anti-AtSGS3 (diluted at 1:3000), anti-AtAGO7 antibodies (diluted at 1:3000), anti-NtSGS3 antibody (diluted at 1:3000) (Yoshikawa et al., 2013), anti-DDDDK-tag mAb (diluted at 1:5000) (Medical & Biological Laboratories) and anti-HA-tag mAb (diluted at 1:5000) (Medical & Biological Laboratories) were used as primary antibodies. Peroxidase AffiniPure Goat Anti-Rabbit IgG (H+L) (diluted at 1:20000) (Jackson ImmunoResearch), Anti-IgG (H+L) (Mouse) pAb-HRP (diluted at 1:5000) (Medical & Biological Laboratories) and Mouse TrueBlot ULTRA: Anti-Mouse Ig HRP (Rockland Immunochemicals, Inc.) (1:1000) were used as secondary antibodies.

### Northern blotting

For *in vitro* assays, two microliter of reaction mixture was mixed with 8 µl of low salt PK solution [0.125% SDS, 12.5 mM EDTA, 12.5 mM HEPES-KOH (pH7.4) and 12.5% Proteinase K (TaKaRa)], and incubated at 50°C for 10 min. Ten microliter of 2 × formamide dye [10 mM EDTA, pH 8.0, 98% (w/v) deionized formamide, 0.025% (w/v) xylene cyanol and 0.025% (w/v) bromophenol blue] was added into the mixture, and further incubated at 65°C for 10 minutes. For *in vivo* assays, total RNA was purified with Trizol reagent (Thermo Fisher Scientific), and 10 µl of 300–500 ng/ul total RNAs were mixed with equal volume 2 × formamide dye. Ten µl of samples were run on a denaturing 1% agarose gel, transferred to the Hybond N+ membrane with capillary blotting and fixed with UV crosslinker. For small RNAs, 10 µl of samples were run on a denaturing 18% acrylamide gel. RNAs were transferred to Hybond N membrane with electro blotting and chemically crosslinked (Pall and Hamilton, 2008). TAS3 variants were detected with Digoxigenin (DIG)-labeled long TAS3 probe (Figure 2D) or 5′ ^32^P-radiolabeled oligo probe mixtures (oligo1230-1234) (Figure 5F). F-TAS1a and its 5′ cleaved fragment were detected with a 5′ ^32^P-radiolabeled oligo probe (oligoA4) (Figure 4E). U6 RNA, miR173, miR390, and tasiRNAs from TAS3 variants were detected with 5′ ^32^P-radiolabeled oligo1129, oligo1353, oligo1131, oligoD7, respectively.

### Immunoprecipitation with anti-FLAG antibody

F-AGO7-RISC or F-AGO1-RISC was assembled as shown above. Target RNAs were mixed with the RISCs at a final concentration of 50 nM, and incubated for 20 min. The reaction mixture was incubated with Dynabeads protein G (Invitrogen) coated with anti-FLAG antibody on a rotator at 4°C for 1 h. The beads were washed three times with 1 × lysis buffer containing 200 mM NaCl and 1% Triton-X 100 or 1 × wash buffer (20 mM Hepes, pH 7.5, 120 mM KCl, 10 mM MgCl2 and 0.2% Nonidet P-40). After removing buffer completely, 1×SDS-PAGE sample buffer was added to the beads. The samples (input, supernatant, and beads) were heated for 5 min and used for SDS-PAGE. Western blotting was performed as described above.

### Immunodepletion of endogenous SGS3 protein

Fifty microliter of BY-2 lysate was mixed with 1.66 µg of anti-NtSGS3 (Yoshikawa et al., 2013) or Normal Rabbit IgG (Medical & Biological Laboratories) at 4°C for 1h. To remove the antibodies and binding proteins thereof, the lysate was mixed with the pellet of 50 µl Dynabeads protein G, and incubated at 4°C for 1h. The supernatant was transferred into new tubes. After flash freezing by liquid nitrogen, the SGS3 or Mock-depleted lysate was stored at -80°C .

### Photoactivated UV crosslinking

*In vitro* reaction mixtures were prepared as outlined above (*Immunoprecipitation with anti-FLAG antibody*) with F-AGO7, ^32^P-labeled miR390_21_4SU, and TAS3-G21A-3M. The sample was transferred to Terasaki plate wells (7 µl/well) and exposed to > 300 nm UV radiation for 15 s using a UV crosslinker (SP-11 SPOT CURE, USHIO) with a uniform radiation lens (USHIO) and a long-path filter (300 nm, ASAHI SPECTRA) at 3 cm from the light. For input sample, aliquots of reaction mixture were transferred into a new tube, and mixed with 4×SDS-PAGE sample buffer. For FLAG-IP, the reaction mixture was incubated with Dynabeads protein G coated with anti-FLAG antibody on a rotator at 4°C for 1 h. For SGS3-IP, the reaction mixture was first incubated with anti- NtSGS3 antibody at 4°C for 1 h, then with Dynabeads protein G at 4°C for another 1 h. The tube was then placed on a magnetic stand to transfer the supernatant into a new tube, which was then mixed with 4×SDS-PAGE sample buffer. The beads were washed three times with 1×lysis buffer containing 800 mM NaCl and 1% Triton-X 100. After removing the buffer completely, 1×SDS-PAGE sample buffer was added to the beads. The samples (input, supernatant, and beads) were heated for 5 min and used for SDS-PAGE. After drying, the gel was exposed to a phosphor imaging plate.

### Agrobacterium-based transient expression in *Nicotiana benthamiana*

The *Nicotiana benthamiana* infiltration assay was performed as previously described (Llave et al., 2000). Briefly, pAT006 and pMDC-plasmids were introduced into *Agrobacterium tumefaciens* GV3101 (pMP90). The *Agrobacterium* cells transformed with TAS3 constructs, AGO7, and miR390 or empty vector (pAT006) were pooled at a ratio of 1:1:2 (total optical density at 600 nm (OD600)) = 1.0). The leaves were harvested at ∼48 h post-infiltration. Total RNA was extracted using Trizol reagent (Thermo Fisher Scientific).

## Data availability

All sequencing data are publicly available in DDBJ, under the accession number DRA010034 (currently undisclosed). All other data are available from the authors upon reasonable request.

## Acknowledgements

We thank James Carrington for providing *Nicotiana benthamiana* seed, *Agrobacterium tumefaciens* GV3101, pMDC32, pMDC32-3×HA-AGO7, pMDC32-TAS3a and a detailed protocol for *Agrobacterium* infiltration, Yukio Kurihara for *Arabidopsis thaliana* (Col-0) seeds, Koreaki Ito and Yuhei Chadani for helpful advice on neutral pH gel electrophoresis analyses, Yuichi Shichino and Mari Mito for technical assistance on ribosome profiling, Keisuke Shoji for kind advice and assistance on NGS data analyses, and Kyungmin Baeg and Yayoi Endo for plasmid construction. We also thank all the members of the Tomari laboratory for discussion and critical comments on the manuscript. We also thank Life Science Editors for editorial assistance. This work was supported in part by JST, PRESTO (grant JPMJPR18K2 to H.-o.I.), Grant-in-Aid for Scientific Research on Innovative Areas (‘Nascent-chain Biology’) (grant 26116003 to H.-o.I.), and Grant-in-Aid for Scientific Research (B) (grant 18H02380 to M.Y.). DNA libraries were sequenced by the Vincent J. Coates Genomics Sequencing Laboratory at UC Berkeley, supported by an NIH S10 OD018174 Instrumentation Grant.

## Author Contributions

H.-o.I. conceived of the project and designed the experiments; H.-o.I. and T.F. performed ribosome profiling and bioinformatic analyses with the supervision of S.I; H.-o.I., A.L., and K.K. performed biochemical analyses; A.M. and A.T. performed transient expression assays in *Nicotiana benthamiana*; H.-o.I., S.I. and Y.T. wrote the manuscript with editing from all the authors; all the authors discussed the results and approved the manuscript.

---

**Figure S1.**
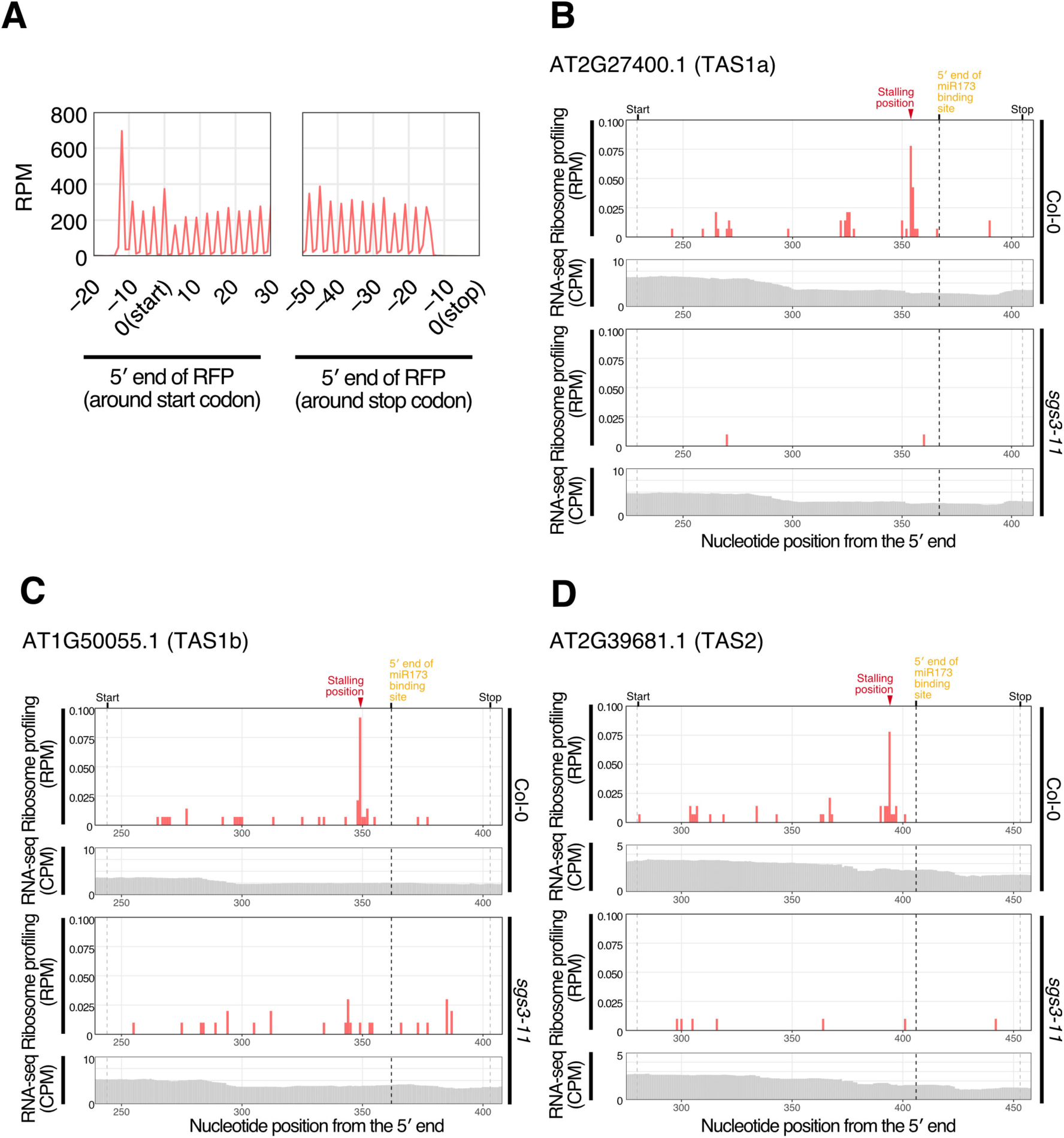
Representative ribosome stalling positions with a downstream miRNA-binding site. (A) Ribosome occupancies around start (left) and stop (right) codons using 28 nt foot prints for 3 day old seedlings of wild-type *Arabidopsis thaliana* (Col-0). The traces indicate 5′ end of ribosome footprints. Ribosome footprints (A-site positions) in RPM and RNA-seq in CPM in 3 day old wild-type or *sgs3-11* mutant seedlings are shown for the following transcripts: (B) AT2G27400.1 (TAS1a), encoding one of the isoforms of TAS1; (C) AT1G50055.1 (TAS1b) encoding one of the isoforms of TAS1; (D) AT2G39681.1 (TAS2) encoding a precursor of tasiRNAs with a miR173 binding site. Related to Figure 1B and C.

**Figure S2.**
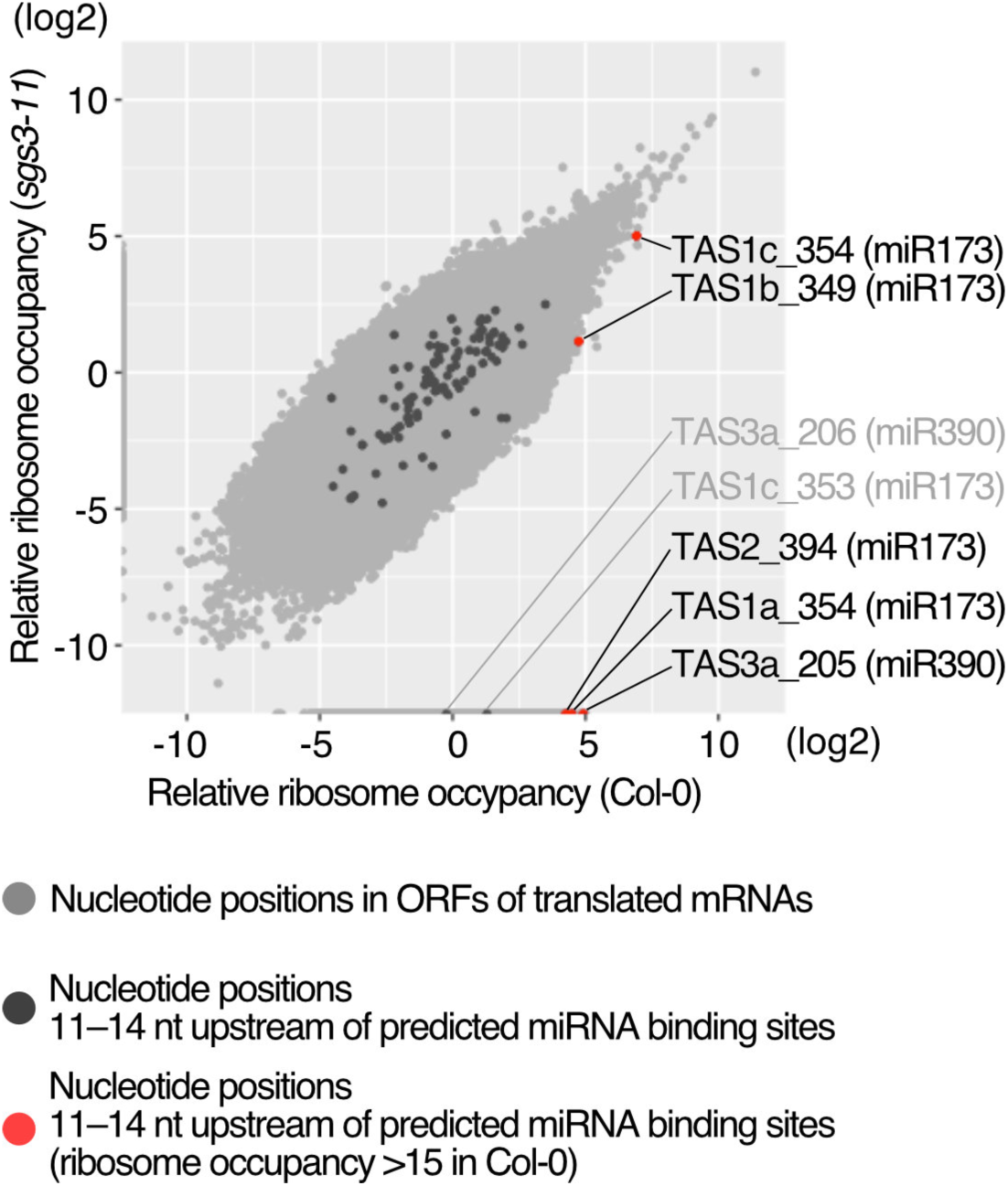
SGS3 is not a general ribosome stalling factor, but rather a specific stalling enhancer for miRNA-mediated ribosome stalling. A scatter plot shows the relative ribosome occupancy (Materials and Methods) between Col-0 and *sgs3-11* seedlings. The nucleotide positions with ribosome footprints (RPM over 0.05 in Col-0 or *sgs3-11*) in translating ORFs are shown in light gray. See also the legend of Figure 1A.

**Figure S3.**
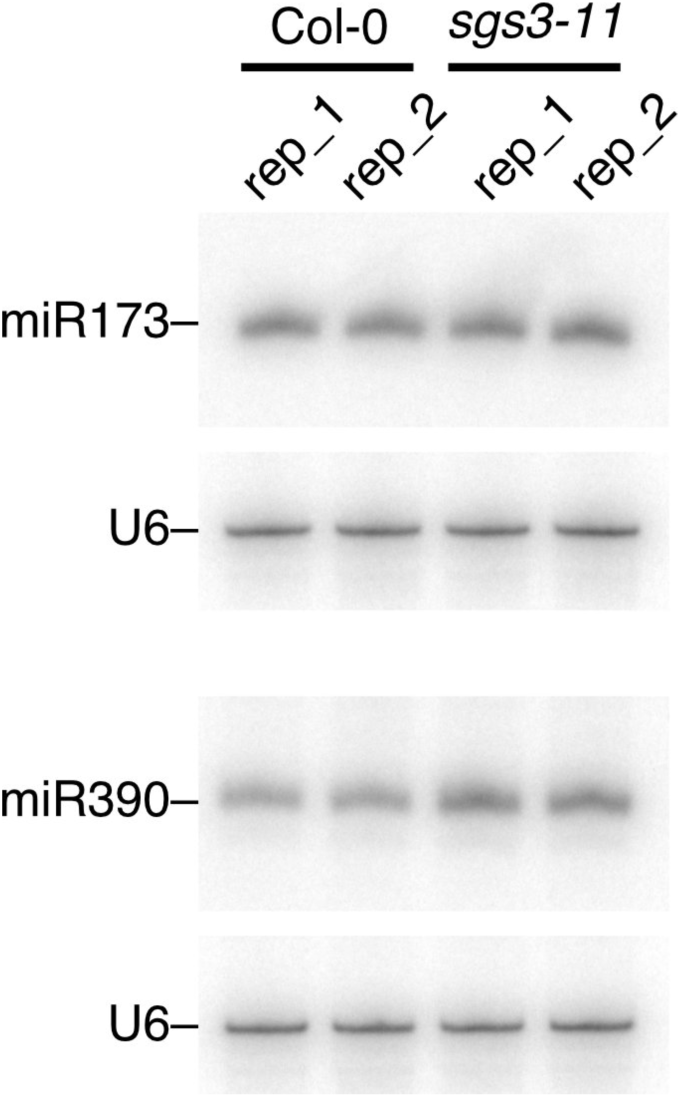
miR173 and miR390 abundance in Col-0 and *sgs3-11* seedlings. miR173 and miR390 in wild-type (Col-0) and *sgs3-11* seedlings were detected by Northern blotting. U6 RNA was used as a loading control.

**Figure S4.**
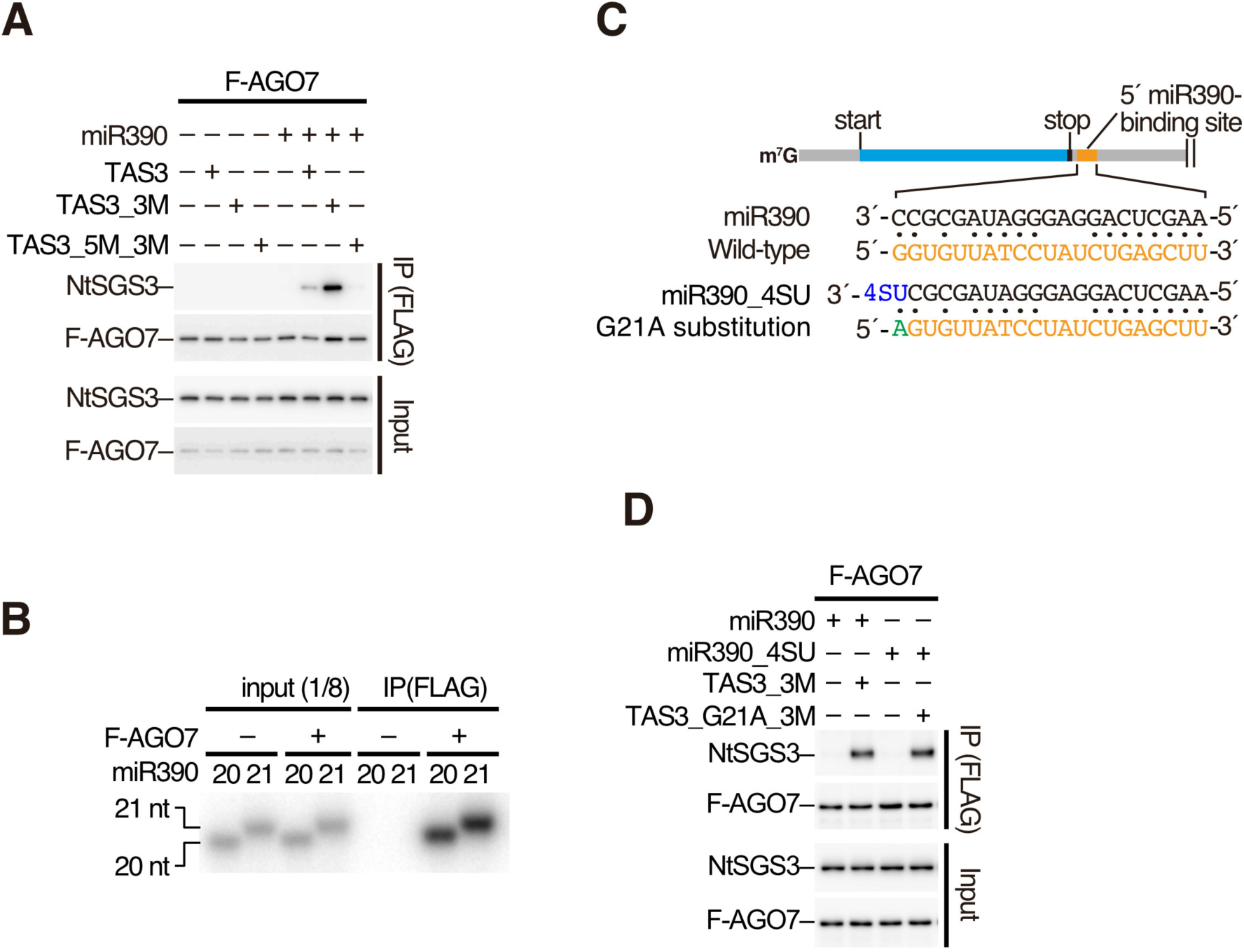
Co-immunoprecipitation of NtSGS3 with AGO7 in the presence of TAS3 variants, and AGO7-RISC assembly with 21-nt and 20-nt miR390. (A) NtSGS3 was specifically co-immunoprecipitated with F-AGO7 in the presence of miR390 duplex and TAS3 or the TAS3-3M. The reason why more SGS3 was co-immunoprecipitated in TAS3_3M than wild-type TAS3 is because the wild-type TAS3 mRNA is cleaved at the 3′ binding site by AGO7-RISC, thereby destabilized in the lysate as shown in Figure 2D. (B) *In vitro* RISC assembly with F-AGO7 and radiolabeled 20 and 21-nt miR390 duplexes. After RISC assembly, F-AGO7 was immunoprecipitated with anti-FLAG antibody. The co-immunoprecipitated miR390 was analyzed by denaturing PAGE. Both 20- and 21-nt miR390 duplexes were efficiently incorporated into AGO7. (C) Schematic of base-pairing configurations between miR390-4SU and a 5′ miR390-binding site with a G21A substitution. The mutated nucleotides in TAS3 variant are shown in green. 4- thiouridine is shown in blue. (D) AGO7-RISC loaded with 21-nt miR390 variant possessing 4- thiouridine at the 3′ end (miR390_4SU) efficiently interacts with NtSGS3 in the presence of TAS3 variant with a compensatory G-to-A mutation at the 5′ end of miR390 binding site (TAS3_G21A_3M).

**Figure S5.**
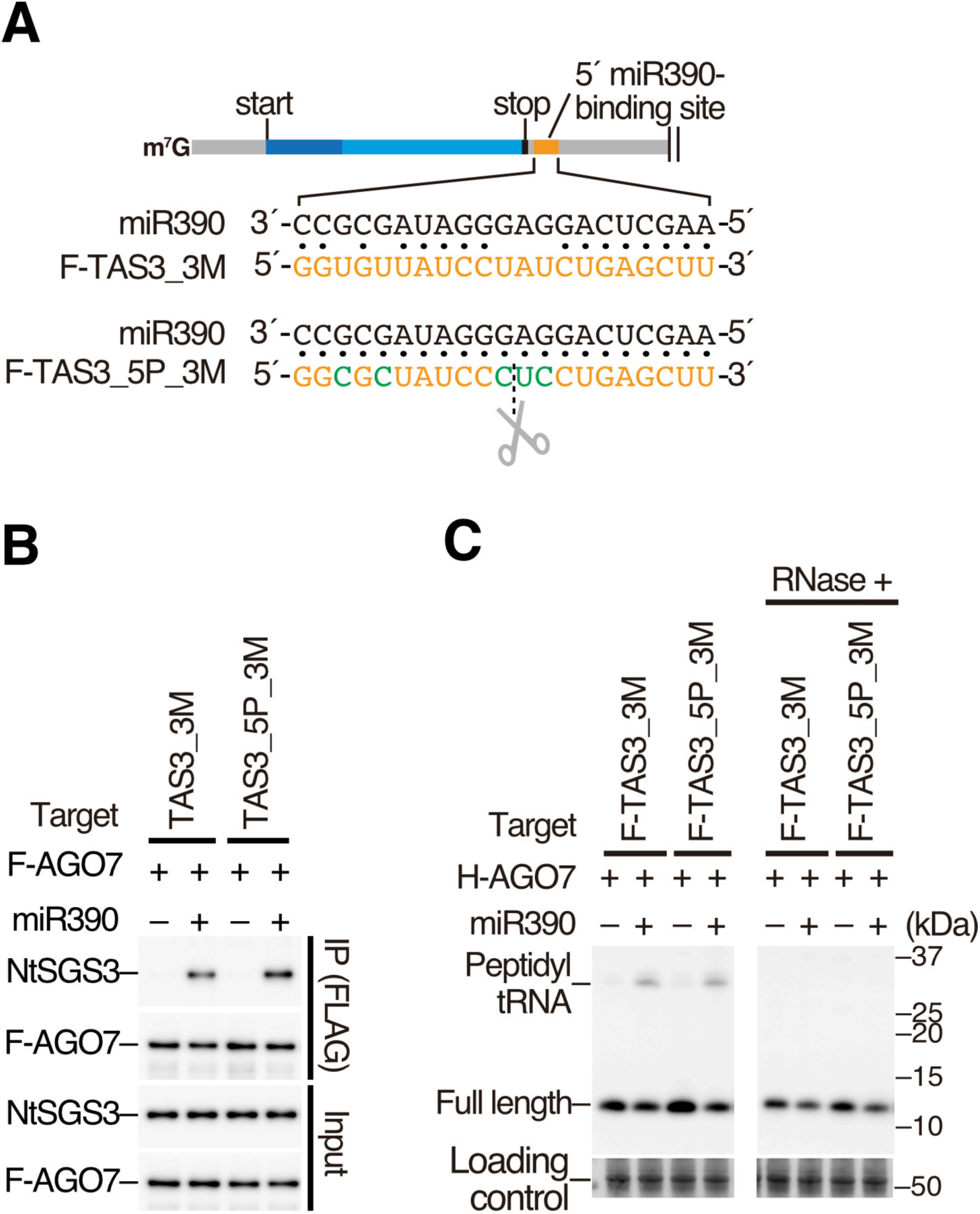
A cleavable target site facilitates ribosome stalling mediated by the miR390-AGO7 RISC. (A) Schematic of base-pairing configurations between miR390 and a 5′ target site with perfect complementarity to miR390. The mutated nucleotides in the TAS3 variant (F-TAS3_5P_3M) are shown in green. (B) Co-immunoprecipitation experiments. AGO7-RISC efficiently interacts with SGS3 in the presence of a F-TAS3_3M variant with a 5′ target site with perfect complementarity to miR390 (F-TAS3_5P_3M). (C) *In vitro* ribosome stalling experiments. Peptidyl-tRNA was accumulated in the presence of AGO7-RISC and F-TAS3_5P_3M, suggesting that cleavable site facilitates ribosome stalling mediated by miR390-AGO7-RISC.

**Figure S6.**
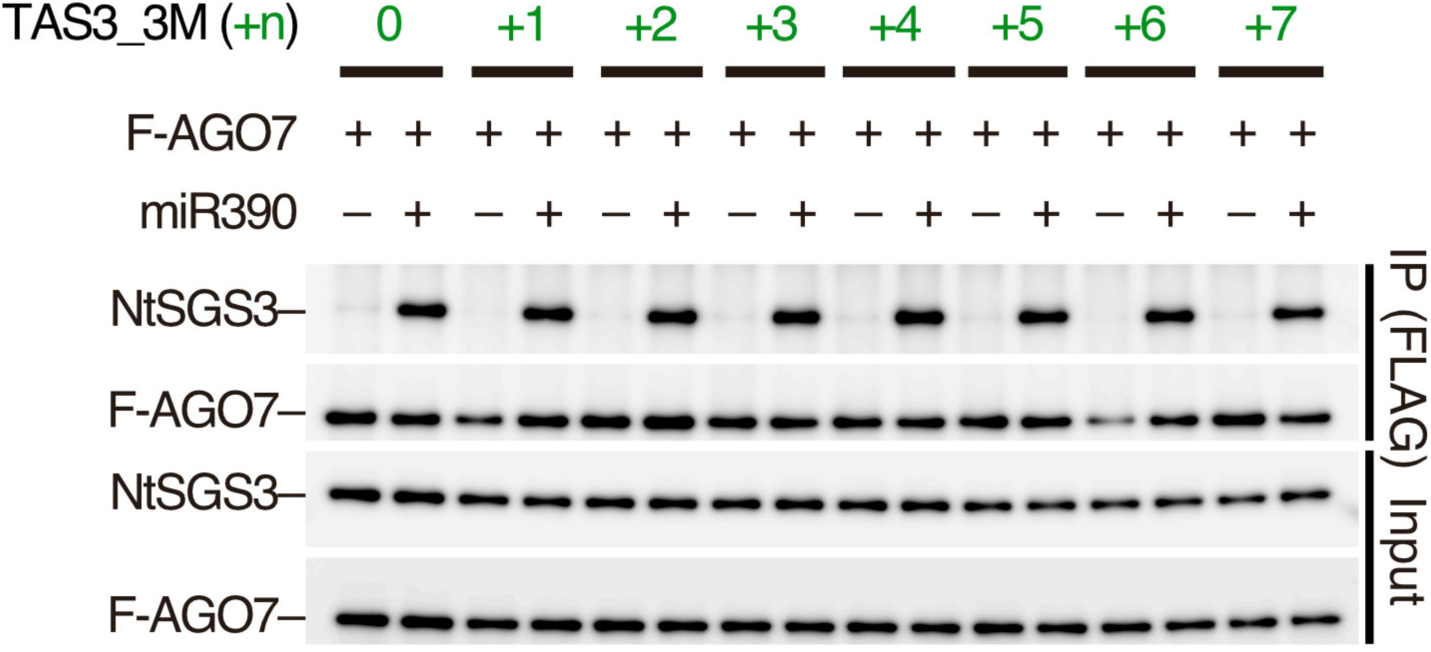
Nucleotide insertion between the stop codon and the 5′ miR390 binding site has no effect on the interaction between AGO7 and NtSGS3. NtSGS3 was specifically and efficiently co-immunoprecipitated with F-AGO7 in the presence of miR390 duplex and TAS3 variants with nucleotide insertions between the stop codon and the 5′ miR390-binding site.

**Table S1.**
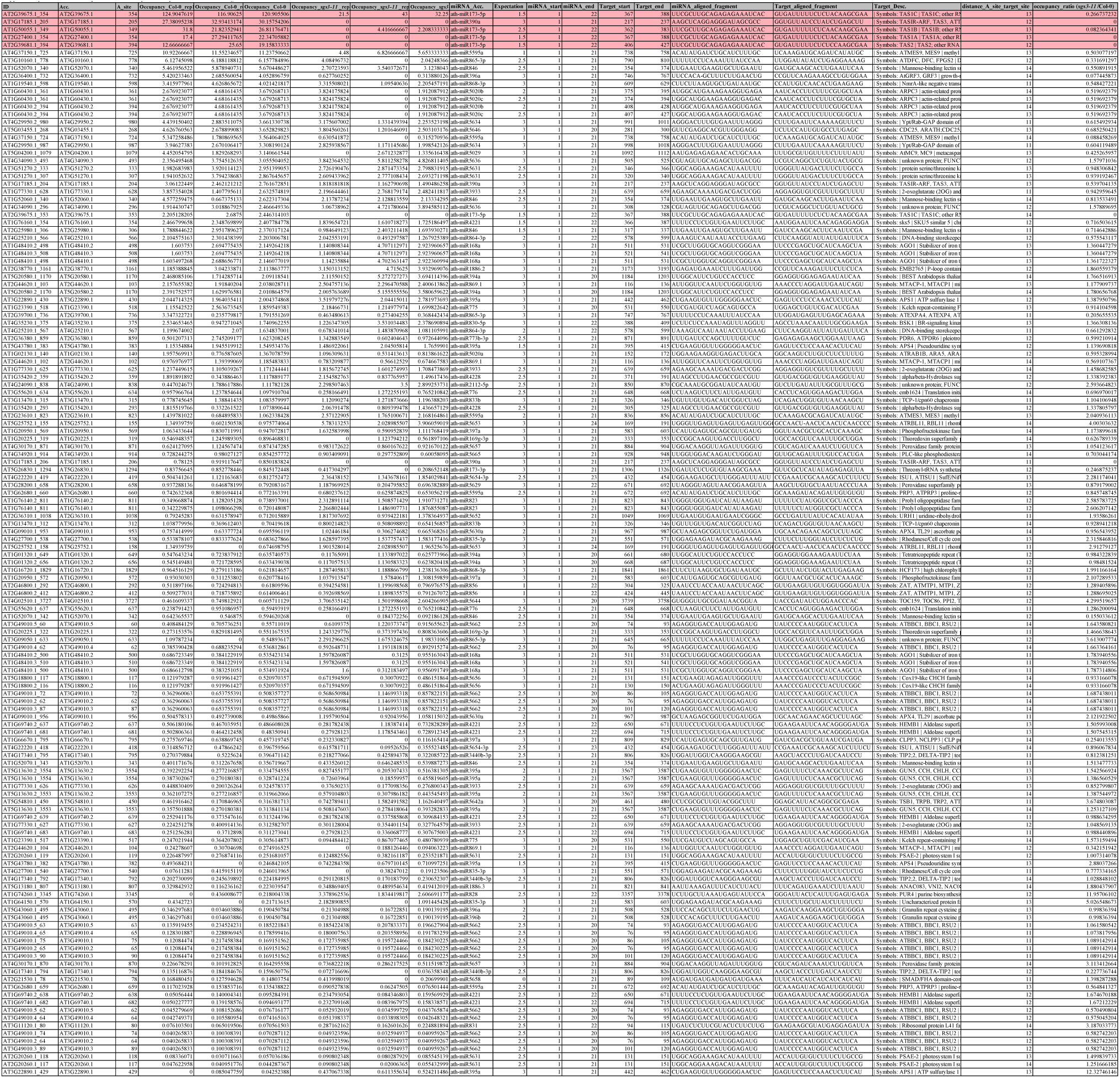
Nucleotide positions 11–14 nt upstream of predicted miRNA binding sites.

**Table S2.**
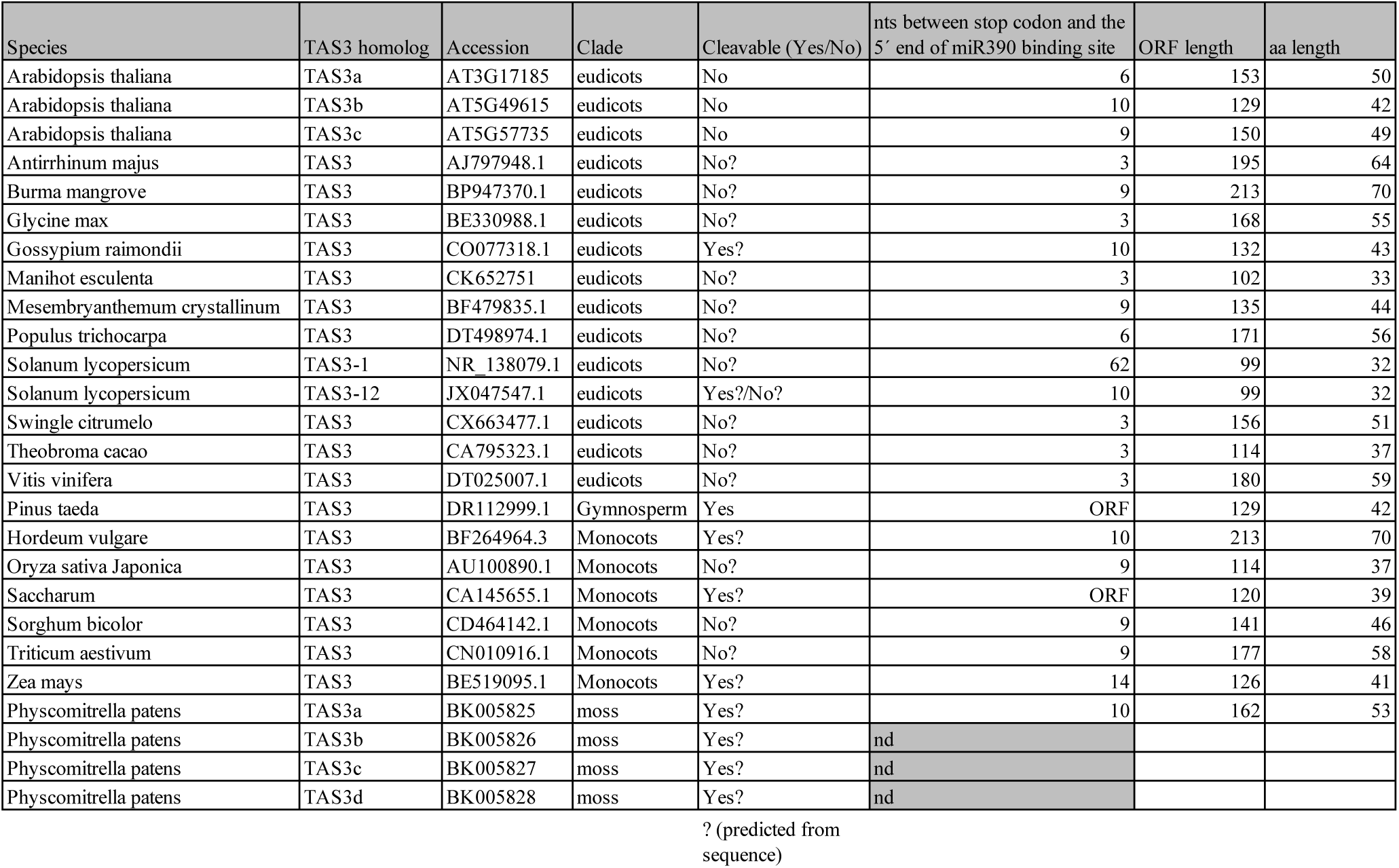
List of TAS3 homologs.

**Table S3.**
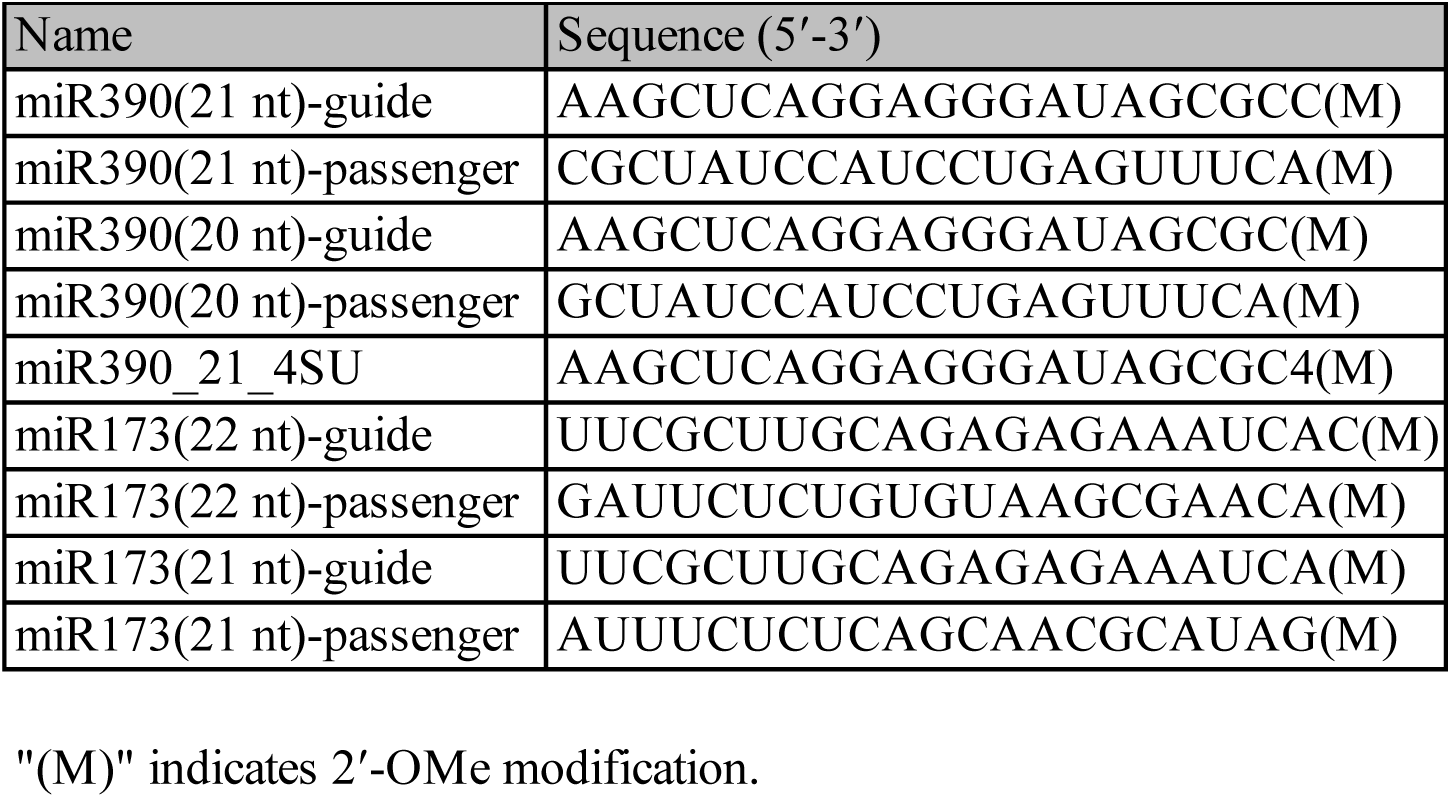
List of synthetic RNA oligos used in this study.

**Table S4.**
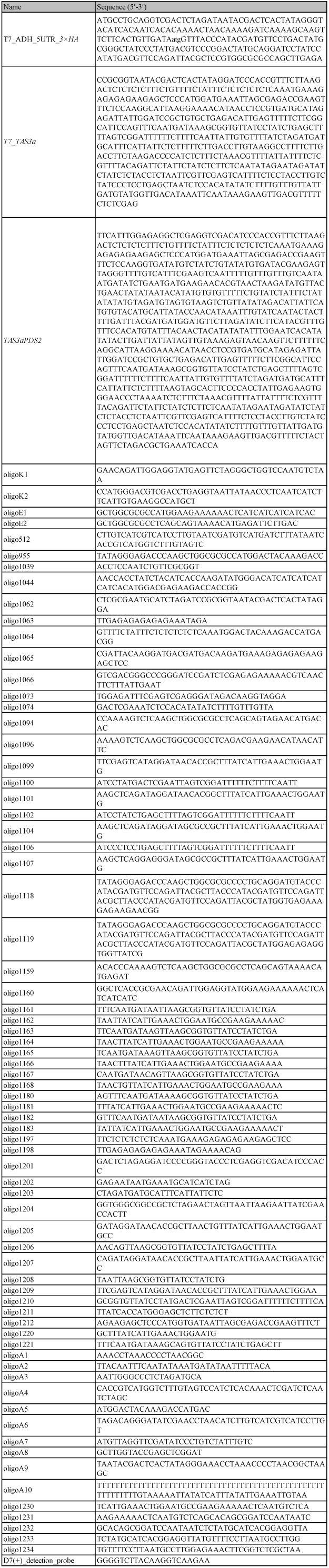
List of synthetic DNA oligos and long DNA fragments used in this study.

